# PHENOTYPIC AND GENETIC CORRELATION BETWEEN SLEEP, BEHAVIOR, AND MACROSCALE CORTICAL GREY MATTER

**DOI:** 10.1101/772681

**Authors:** Masoud Tahmasian, Fateme Samea, Habibolah Khazaie, Mojtaba Zarei, Shahrzad Kharabian, Felix Hoffstaedter, Julia A. Camilleri, Peter Kochunov, B.T. Thomas Yeo, Simon B. Eickhoff, Sofie L. Valk

## Abstract

Humans need about 7 to 9 hours of sleep per night. Sleep habits are heritable, associated with brain function and structure, and intrinsically related to well-being, mental and physical health. This raises the question whether associations between sleep, mental and physical health can be attributed to a shared macroscale neurobiology.

Combining neuroimaging and behavioral genetic approaches in two independent large-scale datasets (n=1887) we demonstrate phenotypic and genetic correspondence between sleep, intelligence, and BMI. Sleep was associated with local thickness variation in frontal, temporal, and occipital cortices. Using a comprehensive multivariate approach, we identified two robust latent components highlighting the interdigitation of sleep, intelligence, BMI, and depression and their shared relation to regions in unimodal and heteromodal association cortices. Latent relationships were heritable and driven by shared additive genetic factors. These observations provide a system-level perspective on the interrelation of sleep, mental, and physical conditions, anchored in grey-matter neuroanatomy.

---

Sleep plays an active role in providing adaptive physiological function^1^, consolidating and retaining new memories^2^, metabolite clearance^3^, hormones’ secretion^4^, and synaptic hemostasis^5^. The National Sleep Foundation suggests seven to nine hours of sleep per night for young adults^6^. However, people in modern societies are suffering from inadequate sleep and its consequences^6^. Sleep loss is associated with impairment in cognitive performance, motor vehicle accidents, and poor quality of life^7, 8^, and contributes to heightened socioeconomic burden^9, 10^. Beyond the quantity of sleep (sleep duration), quality of sleep includes sleep onset latency (i.e., time between going to bed and falling asleep), sleep efficiency (i.e., the percentage of time in bed, during which someone is asleep), sleep disturbances, use of sleeping medication, and daytime dysfunction, all interacting with individual health and well-being^11, 12^. For example, it has been revealed that poor sleep quality is associated with higher rate of depressive symptoms in healthy subjects^13, 14^ and sleep disturbances are common in mood (e.g., depression) and cognitive disorders^15, 16^.

Individual differences in sleep behaviors are heritable^17–19^ and various genetic, metabolic, behavioral, and psychological risk factors have been suggested for the development and maintenance of poor sleep quality and sleep disorders^20–22^. For example, genome wide association studies (GWAS) have associated insomnia disorder to structure of the striatum, hypothalamus, and claustrum, where gene expression profiles show association with the genetic risk profile of such individuals^23, 24^. Moreover, it has been demonstrated that sleep disturbance is associated with hypertension, diabetes, and obesity^25, 26^ whereas depressive symptoms, physical illness, and fatigue were reported as risk factors for both poor sleep quality and short sleep duration^27, 28^. Sleep quantity and quality also both relate to cognitive and academic performance^7, 8^. In addition, various studies have reported sleep disturbance to be associated with alterations in brain structure and function. For example, neuroimaging meta-analyses have implicated structural and functional abnormalities in hippocampus, amygdalae, and insula in patients with sleep apnea^29^ and also indicated convergent functional brain alterations in inferior parietal cortex and superior parietal lobule following acute sleep deprivation^30^. Moreover, white matter integrity underlying prefrontal areas has been associated with sleep duration and sleep quality^31^. Lastly, lower prefrontal gray matter volume has been associated with greater sleep fragmentation in older individuals^32^.

Importantly, it has been demonstrated that macroscale grey matter neuroanatomy is heritable ^33–35^, indicating part of the variance in brain structure can be related to additive genetic effects. Indeed, genetic factors influence cortical thickness in a systematic fashion where both functional and geometric constraints influence genetic correlation between and within brain systems ^36, 37^. Recent studies have indicated that phenotypic correlation between cortical thickness and intelligence, as well as BMI is driven by additive genetic factors^38–40^, suggesting a shared genetic basis of cortical thickness and non-brain traits. This raises the question whether the interrelation of sleep, mental, and physical health could be linked to shared neurobiological mechanisms.

To answer this question, we combined structural neuroimaging approaches in two independent samples, including Human Connectome Project (HCP unrelated sample n=424) as well as the enhanced NKI Rockland sample (eNKI: n=783). The HCP sample consists of young adults only, whereas the eNKI sample consists of both children, younger and older adults, enabling us to evaluate sleep-brain relationship in various age ranges. We conducted phenotypic observations with behavioral genetic approaches in the twin-based full HCP sample (n=1105). Sleep variation was assessed using the Pittsburg Sleep Quality Index (PSQI) ^11^, a widely used questionnaire summarizing self-reported indices of sleep. Our main measures of interest were sleep quantity (self-reported sleep duration), as well as global sleep quality (total PSQI) score, as previous work has associated these factors with brain structure^41, 42^ and genetic variation^43^. Based on previous literature^7, 8, 25–28^ as well as phenotypic correlations in the HCP sample we selected BMI, intelligence, and depression score, to evaluate potential shared neuroanatomical basis between sleep and mental and physical aptitudes. In the HCP sample intelligence is summarized as Total Cognitive Score, based on the NIH Toolbox Cognition^44^, whereas in the eNKI sample intelligence is measured using the Wechsler Abbreviated Scale of Intelligence (WASI-II)^45^. Depression was measured using the ASR depression DSM-oriented scale for Ages 18-59^46^ in the HCP sample, and using Beck Depression Inventory (BDI – II) in the eNKI sample. BMI was calculated at weight/squared(height) in both datasets.

Our analysis revealed a phenotypic relationship between sleep and depression, BMI, and intelligence in both HCP sub-sample including only unrelated individuals as well as the eNKI sample. Following, we demonstrated depression, BMI, and intelligence were heritable and observed genetic correlation between sleep quantity and quality, BMI, and intelligence in the twin-based HCP sample, indicating sleep hygiene displays pleiotropy with these factors in the mentioned samples. Analysis of heritability and co-heritability was performed with maximum likelihood variance-decomposition methods using Sequential Oligogenic Linkage Analysis Routines (www.solar-eclipse-genetics.org; Solar Eclipse 8.4.0.) Heritability (h^2^) is the total additive genetic variance and genetic (ρg) correlations were estimated using bivariate polygenic analyses. Using an atlas-based approach to summarize cortical thickness^47^, we observed significant local associations between sleep duration and cortical structure in both samples which were driven by additive genetic factors. Post-hoc analysis indicated that variance in intelligence and BMI also related to thickness in areas associated with sleep duration. Subsequently, based on our observation that sleep relates to BMI, intelligence, and depression as well as to cortical thickness, we performed partial least squares (PLS) analysis, to identify latent relationships between these factors. PLS is a multivariate data-driven approach, enabling simultaneous linking of behavioral measures to brain structure. We identified two significant and robust latent factors spanning distinct neurocognitive dimensions. Using the twin-structure of HCP we observed these factors are heritable and their relation driven by shared genetic effects. Taken together, the current study highlights the interrelation of sleep, mental and physical health, which is reflected by shared neurobiological signatures.

## Results

### Data samples

We studied two independent samples from openly-shared neuroimaging repositories, Human Connectome Project and enhanced eNKI. Human Connectome Project (HCP; http://www.humanconnectome.org/), comprised data from 1105 individuals (599 females), 285 MZ twins, 170 DZ twins, and 650 singletons, with mean age 28.8 years (SD = 3.7, range = 22–37). For phenotypic analysis we selected unrelated individuals resulting in a sample of 424 (228 females) individuals with mean age of 28.6 years (SD =3.7, range =22-36). Our second sample was based on the Nathan Kline Institute-Rockland Sample (eNKI), made available by the Nathan-Kline Institute (NKY, NY, USA), (https://www.ncbi.nlm.nih.gov/pmc/articles/PMC3472598/). This sample consisted of 783 (487 females) individuals with mean age of 41.2 years (SD =20.3, range =12-85) enabling us to identify life-span relations between sleep, brain structure and behaviour. Details on the sample characteristics can be found in the Methods section.

### Relation between sleep, mental and physical health

First, we sought to evaluate whether our measures of mental and physical status related to sleep quantity and quality. Here we correlated sleep duration and global sleep quality to phenotypic variation in cognition, mental, and physical health (for selection of markers please see Table S1). This data-driven analysis in the HCP phenotypic data revealed that cognitive, mental and physical phenotypic variation have a strong relation to variation in sleep (Table S2 and Table S3). Given the marked role of both depression as well as BMI on both sleep duration and global sleep quality, we selected these as phenotypes of interest for further analysis. As we observed many cognitive factors related to sleep duration and sleep quality, we selected general intelligence, as this marker has been shown to be highly heritable and related to brain structure^39^ Next, we successfully evaluated that depression, IQ, and BMI showed moderate phenotypic inter-correlations in unrelated HCP and eNKI samples as well as full HCP-sample (Table S4). Evaluating the relation between sleep and our selected markers in eNKI, in addition to HCP, we observed that sleep duration had a consistent negative phenotypic relation to both BMI and depression, and a positive relation to IQ. Depression, IQ, and BMI were all significantly heritable (Table 1) and we observed a negative genetic correlation between BMI and IQ (ρg =-0.27 (0.06)). Moreover, sleep duration was significantly heritable (h2=0.24, p<0.001), and phenotypic correlations were mirrored by genetic correlations, were we observed sleep duration to show a positive genetic correlation with IQ (p<0.001), but negative with BMI (p<0.01) (Table 1). Depression only showed an environmental correlation with sleep duration.

**Table 1.** Sleep has neurogenetic overlap with depression, BMI and IQ. Cross-sample analysis of genetic correlational analysis between sleep duration and global sleep quality on the one hand, and depression, BMI, and IQ on the other, including 95% confidence intervals. * indicates p<0.05, ** indicates FDR q<0.05.

| **Sleep duration ($h^2= 0.24\pm0.06$)** |  |  |  |
| --- | --- | --- | --- |
| *Sample* | *Depression*<br>($h^2=0.24\pm0.06$) | *BMI*<br>($h^2=0.68\pm0.04$) | *IQ*<br>($h^2=0.66\pm0.04$) |
| HCP (unrelated sample) | -0.09 [-0.19 0.00] | -0.11 [-0.21 -0.02] * | 0.11 [0.01 0.19] * |
| eNKI | -0.16 [-0.24 -0.07] ** | -0.17 [-0.24 -0.09] ** | 0.11 [0.04 0.18] * |
| HCP (total sample) | -0.07 [-0.13 -0.02] * | -0.14 [-0.19 -0.08] ** | 0.09 [0.03 0.15] * |
| Genetic correlation (HCP) | 0.17(0.20) | -0.33 (0.11) * | 0.42 (0.11) ** |
| Environmental correlation (HCP) | -0.16(0.06) * | 0.01 (0.07) | 0.19 (0.06) * |
| **Global sleep quality ($h^2= 0.12 \pm 0.06$)** |  |  |  |
| *Sample* | *Depression* | *BMI* | *IQ* |
| HCP (unrelated sample) | 0.37 [0.29 0.45] ** | 0.14 [0.04 0.23] * | -0.07 [-0.16 0.03] |
| eNKI | 0.37 [0.29 0.45] ** | 0.09 [0.02 0.17] * | -0.09 [-0.16 -0.02] * |
| HCP (total sample) | 0.35 [0.30 0.40] ** | 0.10 [0.04 0.16] * | -0.10 [-0.16 -0.04] * |
| Genetic correlation (HCP) | 0.32(0.26) | 0.41 (0.16) * | -0.59 (0.20) ** |
| Environmental correlation (HCP) | 0.38(0.05) ** | 0.03 (0.07) | 0.17 (0.06) * |

Global sleep quality showed comparable relations to depression, IQ, and BMI, with strong phenotypic correlation between poor sleep quality (higher total PSQI score) and higher depression and BMI scores, as well as between poor sleep quality and lower IQ across samples (Table 1). Global sleep quality was also influenced by additive genetic effects (h2=0.12, p<0.05), but less so than sleep duration. Phenotypic correlations were paralleled by genetic correlations, were poor sleep quality were genetically correlated with higher IQ (p<0.001) and higher BMI (p<0.01). Again, depression only showed environmental correlation with global sleep quality (p<0.05) (Table 1).

### Phenotypic correlation between sleep and brain structure in two independent samples

Next, we evaluated the phenotypical relation between sleep indices (global sleep quality and sleep duration) and cortical thickness in both the unrelated subsample from the Human Connectome Project (HCP) (n=424) as well as eNKI (n=783). Behaviorally, we observed a strong negative correlation (Spearman r=-0.51 [-0.59 −0.44], p<0.001) between global sleep quality and sleep duration (Fig. 1A). Correlation of sleep indices with brain structure demonstrated a negative link between left superior frontal thickness (area 6d2 and pre-supplementary motor area) and sleep duration (r=-0.1, FDRq<0.02), that remained significant when controlling for self-reported depressive symptoms, as well as intake of sleep medications, intelligence, and BMI. Global sleep quality did not relate to local variations in cortical thickness (Fig. 1B,C). However, when evaluating the relationship in the complete HCP sample including twins and siblings, we observed only a trending relation between sleep duration and cortical thickness (Figure S1).

**Figure 1.**
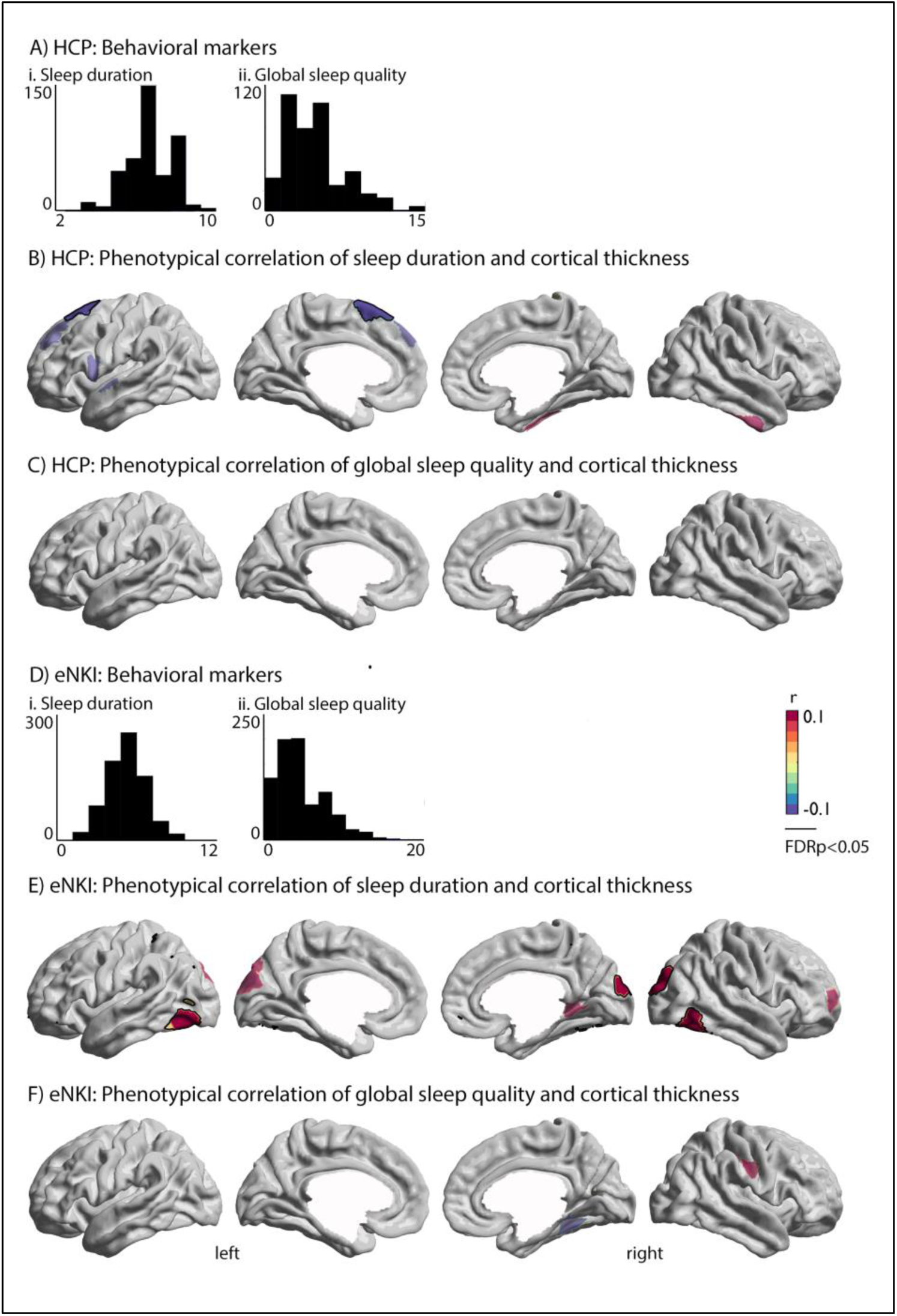
Patterns of phenotypical correlation between sleep duration and cortical thickness in HCP and eNKI samples. A) distribution of variables in the unrelated HCP subsample; B+C) phenotypical correlation of sleep duration/global sleep quality and cortical thickness; D) distribution of variables in the eNKI sample, as well as the correlation between sleep duration and global sleep quality score; E+F) phenotypical correlation of sleep duration/global sleep quality and cortical thickness. Red indicates a positive relationship, whereas blue indicates a negative phonotypical relationship between sleep and brain structure. Whole-brain findings were corrected for multiple comparisons using FDR correction (q<0.05, black outline). Significant associations between sleep indices and brain structure have black outline, whereas trends (p<0.01) were visualized at 60% transparency.

In eNKI, we replicated the negative behavioral correlation between sleep duration and global sleep quality (Spearman r=-0.53 [-0.58 −0.47], p<0.001). Though we again found no relation between global sleep quality and cortical brain structure, sleep duration showed a positive link between bilateral inferior temporal regions (left: 0.13, FDRq<0.02, right: r=0.12 FDRq<0.02) and right occipital cortex (r=0.14, FDRq<0.02). Findings remained significant when controlling for self-reported depressive symptoms, as well as intake of sleep medications, intelligence, and BMI.

### Replication analysis of correspondence between sleep duration and cortical thickness

As we observed divergent local phenotypic correlations between sleep duration and cortical thickness in two independent large samples, we evaluated the inconsistencies across samples more precisely. Indeed, post-hoc analysis indicated that local effects of phenotypic correlations varied strongly in magnitude across samples in phenotypic analysis (Table 2). At the same time, cortex-wide patterns of correlation were similar across datasets and analyses, indicating that the direction of sleep-thickness associations are similar across both samples. Indeed, we observed a high overlap between spatial distribution of phenotypic correlations between sleep duration, but not global sleep quality, and cortical thickness across samples and subsamples (Table S5). This suggests that the relation between sleep and cortical thickness might be robust at the inter-regional level rather than in local effects only. Additionally, we observed that both intelligence and BMI related to local thickness associated with sleep duration (Table S6), suggesting sleep, intelligence and BMI are dependent on overlapping macro-anatomical structures.

**Table 2.** Inconsistency of associations between sleep duration and cortical thickness across samples and analysis. Cross-sample replication of FDR-corrected ROIs from phenotypic correlational analysis in Fig.1. ** indicates to significant correlation (q<0.05) and * indicates association at trend-level p<0.05.

| **Phenotypic correlation, Figure 1A (HCP)** |  |  |
| --- | --- | --- |
| *Left superior frontal gyrus* | r | p |
| HCP (unrelated sample) | -0.19 | 0.00007** |
| eNKI | -0.03 | 0.40 |
| HCP (total sample) | -0.10 | 0.0013* |
| Genetic correlation (HCP) | -0.27 | 0.035* |
| Environmental correlation (HCP) | 0.02 | 0.97 |
| **Phenotypic correlation, Figure 1B (eNKI)** |  |  |
| *Left inferior temporal cortex* | r | p |
| HCP (unrelated sample) | 0.09 | 0.07 |
| eNKI | 0.13 | 0.0001** |
| HCP (total sample) | 0.04 | 0.14 |
| Genetic correlation (HCP) | -0.09 | 0.67 |
| Environmental correlation (HCP) | 0.06 | 0.27 |
| *Right occipital cortex* | r | p |
| HCP (unrelated sample) | 0.07 | 0.13 |
| eNKI | 0.14 | 0.0001** |
| HCP (total sample) | 0.04 | 0.20 |
| Genetic correlation (HCP) | 0.28 | 0.03* |
| Environmental correlation (HCP) | -0.08 | 0.19 |
| *Right inferior temporal cortex* | r | p |
| HCP (unrelated sample) | 0.05 | 0.30 |
| eNKI | 0.12 | 0.0006** |
| HCP (total sample) | 0.004 | 0.90 |
| Genetic correlation (HCP) | 0.38 | 0.02* |
| Environmental correlation (HCP) | -0.11 | 0.06 |

### Phenotypic correlations between sleep and cortical thickness are driven by additive genetic effects

Next, we evaluated whether phenotypic correlations between sleep duration and cortical thickness were mirrored by additive genetic effects using the twin-structure of the HCP dataset. First, we confirmed that cortical thickness was significantly heritable in the current sample (Figure S2, Table S7). Second, we assessed whether phenotypic correlations observed in Figure 1 were driven by shared additive genetic effects. We found that both frontal cortex (based on HCP) as well as right occipital cortex and right inferior temporal cortex (based on eNKI) were had genetic correlation (p<0.05) with sleep duration (Table 3). Using a whole-brain approach we identified a negative genetic correlation between sleep duration and bilateral frontal cortices thickness, mainly in the bilateral superior frontal gyrus and frontal pole, areas p32 and Fp2 (left: ρ_e_=0.12, p<0.06, ρ_g_=-0.46, FDRq<0.025; right: ρ_e_=0.15, p<0.01, ρ_g_=-0.46, FDRq<0.025) (Figure S1). Findings were robust when controlling for intelligence, BMI or depression score (Table S8). However, though we observed genetic correlation to reflect phenotypic correlation when assessing whole-brain associations (Table S5), we could not observe significant or trend level relationships between local frontal thickness and sleep in the eNKI sample (Table S9). Last, though did not observe significant genetic correlation between global sleep quality and brain structure, we identified a significant environmental relation between global sleep quality and left precentral thickness (r=0, ρ_e_=0.22, p<0.0002, ρ_g_=-0.64, p<0.0003) (Figure S3).

**Table 3.** Phenotypic associations between sleep indices and cortical thickness are reflected by genetic correlations. Genetic and environmental correlation between sleep and thickness in FDR-corrected ROIs from phenotypic correlational analysis in Fig.1. * indicates to significant correlation (q<0.05).

| **Phenotypic correlation, Figure 1A (HCP)** |  |  |
| --- | --- | --- |
| *Left superior frontal gyrus* | r | p |
| HCP (total sample) | -0.10 | 0.0013* |
| Genetic correlation (HCP) | -0.27 | 0.035* |
| Environmental correlation (HCP) | 0.02 | 0.97 |
| **Phenotypic correlation, Figure 1B (eNKI)** |  |  |
| *Left inferior temporal cortex* | r | p |
| HCP (total sample) | 0.04 | 0.14 |
| Genetic correlation (HCP) | -0.09 | 0.67 |
| Environmental correlation (HCP) | 0.06 | 0.27 |
| *Right occipital cortex* | r | p |
| HCP (total sample) | 0.04 | 0.20 |
| Genetic correlation (HCP) | 0.28 | 0.03* |
| Environmental correlation (HCP) | -0.08 | 0.19 |
| *Right inferior temporal cortex* | r | p |
| HCP (total sample) | 0.004 | 0.90 |
| Genetic correlation (HCP) | 0.38 | 0.02* |
| Environmental correlation (HCP) | -0.11 | 0.06 |

### Latent relation between sleep, brain and behavior

As we observed 1) phenotypic and 2) genetic correlations between sleep, intelligence, BMI and, in part, depression, as well as 3) significant but inconsistent relation between sleep duration and cortical thickness, we utilized a multivariate data-driven approach to evaluate latent relationships between sleep, intelligence, BMI and depression on the one hand, and cortical thickness on the other. Indeed, it has been suggested multiple comparison corrections in mass univariate analysis may result in missing effects and thus inconsistencies in the results and a more comprehensive picture of the associations could be gained by a multivariate approach^48^. Here, our primary analysis sample is eNKI, as this enables us to replicate phenotypic and evaluated genetic correlations between latent structures using the full HCP sample. In the eNKI sample we observed two significant latent relationships between our behavioral phenotypes and cortical thickness, controlling for effects of age, sex, and global thickness, explaining 41% of the shared variance (first latent component; p<0.001, association between behavior and brain saliencies: r=0.38), and 25% of the shared variance (second latent component; p<0.01, association between behavior and brain saliencies: r=0.29).

The first component related positively to both sleep duration(r=0.49) and intelligence (r=0.83), and negatively to sleep quality (r=-0.43), BMI (r=-0.46) and depression (r=-0.21). Here intelligence had the strongest effect on the latent relationships. The brain saliency loadings were positively (bootstrap ratio>2) associated sensory-motor areas as well as superior temporal areas, and parahippocampal structures, and negatively with lateral and medial frontal cortex, as well as inferior temporal lobe, precuneus, and posterior parietal cortex. Further qualifying the brain saliency, we observed positive relations were mainly in visual, sensorimotor and limbic functional networks, whereas negative relations were predominantly in the dorsal-attention, fronto-parietal and default-mode networks. Replicating the association in HCP using the behavioral and brain loadings, we identified a significant relation between latent brain and behavioral factors in this sample as well (component 1: r=0.25, p<0.001). Moreover, both brain and behavioral latent factors were significantly heritable (brain saliency: h2±std = 0.63±0.04; behavior saliency: 0.76±0.03) and showed genetic correlation (ρ_e_=-0.07±0.07, p=ns, ρ_g_=0.38±0.05, p<0.0001). The first brain-behavior component seems to reflect a positive-negative axis of behavior, relating high sleep quantity to positive behaviors whereas low quality related negatively to this factor.

The second component related positively to depression (r=0.70), BMI (r=0.50), intelligence (r=0.16), and reduced sleep quality (r=0.69), and negatively to sleep duration (r=-0.43). Positive brain loadings (bootstrap ratio>2) were located in the left sensorimotor areas, right precuneus, and right parietal areas. Negative loadings (bootstrap ratio<-2) were located in left dorsolateral areas, left mid-cingulate, right dorsolateral frontal cortex, and left anterior-mid cingulate. Qualitative analysis revealed positive loading were predominantly in sensorimotor, visual, dorsal attention and default networks, whereas negative loadings were associated with the fronto-parietal, ventral attention, limbic and default networks. Again, we successfully replicated this association in HCP using the behavioral and brain loadings (r=0.10, p<0.002). Both brain and behavioral saliency of the second component were significantly heritable (brain saliency: h2±std = 0.72± 0.03; behavior saliency: 0.51± 0.05) and showed genetic correlation (ρ_e_=-0.11±0.07, p=ns, ρ_g_=0.22±0.07, p<0.0001). This time, sleep quality, depression, BMI, and intelligence showed positive latent relations, but amount had a negative relation to the behavioral saliency, indicating sleep quality has both positive and negative relationships to intelligence. Supplementary analysis in the HCP sample embedded sleep quantity and quality in a broad range of cognitive and emotional behaviors (Table S10 and S11), further highlighting the interdigitation of nocturnal and daytime behaviors.

**Figure 2.**
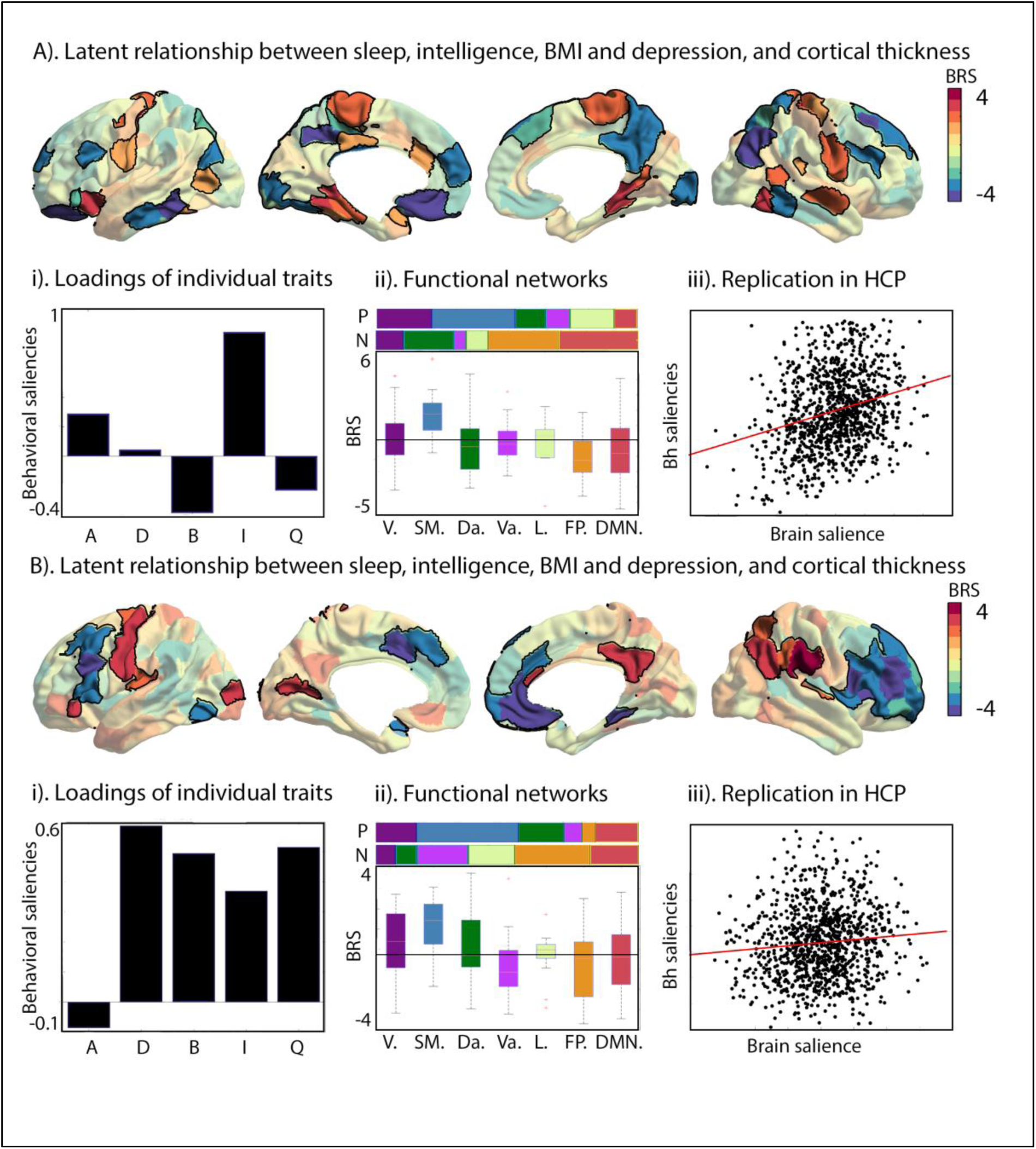
Two latent dimensions of cortical macrostructure and components of sleep, mental and physical health. A). Bootstrap ratio of the first brain saliency that showed significant robustness, where parcel-wise saliencies of BSR>2 are highlighted. Red indicates a positive association whereas blue indicates a negative association; i. Loadings of the individual traits; ii. Relative distribution of positive(P) and negative (N) −2>BSR>2 scores per functional networks^49^, and average BSR in functional networks49 (V=visual, SM=sensorimotor, Da=dorsal-attention, Va=ventral attention, L=limbic, FP=frontopolar, DMN=default mode network), iii. Replication of brain – behavior saliency association in the HCP sample; and B. Relation between brain and behavioral saliencies in HCP sample of the second brain saliency. i. Loadings of the individual traits; ii. Relation to functional networks49 and iii). Relation between brain and behavioral saliencies of second factor in the HCP sample.

## Discussion

Sleep is key to normal human functioning and associated with brain structure and function. At the same time, individual differences in sleeping behavior are heritable and have substantial overlap with cognition, physical, and mental health. This raises the question whether shared variance in sleep, intelligence, BMI, and depression could be due to a shared relationship to macroscale grey-matter anatomy. We combined computational approaches from behavioral genetics and big-data neuroimaging to evaluate the interrelation between sleep, macroscale brain structure, and mental and physical health. Indeed, in two independent samples, we observed that sleep duration as well as global sleep quality had a significant phenotypic relation to intelligence and BMI, which was mirrored by additive genetic effects. Depression showed only phenotypic correlation with sleep. Following, we demonstrated that sleep duration, but not global sleep quality, had significant (but inconsistent) relation with local variance in cortical thickness in both samples. Three out of 4 phenotypic relations between sleep duration and local cortical thickness were driven by additive genetic factors. At the same time, both intelligence and BMI (to a lesser extent) related to variance in cortical thickness in these regions, suggesting these factors might have an overlapping neuroanatomical basis. Consistent with these results, a comprehensive multivariate analysis revealed two robust and heritable signatures highlighting shared relationships between macroscale anatomy and sleep, intelligence, BMI and depression. Both components featured brain structure in both unimodal and heteromodal association areas, and underlined the embedding of nocturnal behavior in daytime functioning. Collectively, our multi-sample approach provides novel evidence that sleep is intrinsically interrelated with macroscale grey matter structure, mental, and physical health.

Our observations highlight the key relation between cognitive abilities, mental and physical health on sleep profile of healthy subjects. Previous work has implicated the role of sleep on life functioning, such as cognitive performance and quality of life^7, 8^, as well as higher rate of depressive symptoms^13, 14^ and hypertension, diabetes, and obesity^25, 26^. Indeed, clear associations of sleep, cognitive performance and behavioral problems have been observed in children^50^, adults^51^, and elderly ^41^. In line with our observations, Lim and Dinges report a relation between complex attention on working memory. In addition, short sleep duration is associated with poor overall IQ measures/cognitive performance in healthy children^52^. BMI and intelligence showed opposite relationship with sleep duration in our study, and IQ score is reported to be correlated with higher BMI in a large sample of 1151 children^53^.

There are various hypotheses on the biological processes underlying the active role of sleep in the neuronal processing of information and consequently mental processing. The trace reactivation or replay hypothesis^54, 55^ suggests that sleep helps memory consolidation through reactivation of traces of neuronal activity patterns encoding information. The synaptic homeostasis hypothesis proposes that sleep is necessary to counterbalance the increase of synaptic connectivity^5^. Last, converging evidence suggests a role of sleep in maintaining functional integrity of the frontoparietal networks that support sustained attention^56, 57^ as well as default mode network^58^, two brain networks implicated in task-unrelated thought. Indeed, in our multivariate analysis we observed a shared relation of intelligence and sleep with cortical thickness in these networks.

At the same time, our work shows that inadequate sleep is implicated heightened BMI. It has been shown that high BMI associated with the abnormal sleep duration and vice versa^59^. Short term sleep restriction is associated with impaired glucose metabolism, dysregulation of appetite and increased blood pressure, and prospective studies found increased risk of weight gain associated with inadequate sleep ^60, 61^. In the same vein, various lines of evidence have related also BMI to brain structure and function^38^, suggestive of a bidirectional relation between sleep, BMI, and brain structure.

Last, we observed a relation between sleep and depressive symptoms. A recent meta-analysis on prospective studies implicated both long and short sleep to be associated with increased risk of depression in adults ^62^. Though the mechanisms underlying this association are not fully understood, daytime tiredness, resulting in increased negative events and emotions, has been shown to be predictive of poor outcome of depression. Next to this, sleep abnormalities relate also low physical activity, which in turn modulates risk of depression. Importantly, sleep factors can predispose, precipitate, and perpetuate depression and in our multivariate model we observed both neutral and positive associations between depression and negative sleep behaviors, highlighting the complex relation between sleep and mental health.

Though we could establish phenotypic as well as genetic correlations between sleep duration and local cortical thickness in two independent samples, findings were inconsistent. In the HCP sample, sleep duration was linked to thickness in frontal areas. The important role of frontal cortex in sleep is previously well-documented. For example, sleep deprivation influences frontal executive functions in both health individuals and patients ^63 64 65^. In addition, sleep deprivation disrupts leads to lower metabolism in the frontal cortex, while sleep recovery moderately restores frontal lobe functions^66^. Function abnormalities are also mirrored by abnormalities in macro-anatomical structure, where cortical thinning in bilateral precentral cortex and the superior/mid frontal cortex related to lower sleep quality^67^ and insomniac patients showed grey matter abnormalities in frontal cortices^68 69^. Here, phenotypical analyses in the eNKI sample demonstrated that sleep duration had a positive link with bilateral inferior temporal regions and right occipital cortex. Indeed, also function and structure temporal and occipital areas have been associated in sleep patterns. For instance, older adults with short or long sleep duration had higher rates of cortical thinning in the frontal and temporal regions, as well as the inferior occipital gyrus^70^ and insomnia has been related to functional abnormalities not only on frontal regions, but also in temporal and occipital areas^71 72^. These excessive hyperarousal, impaired alertness, auditory-and vision-related inattention and experiencing negative moods in such patients. Thus, local effects of sleep on brain structure and function are observed both in the literature as well as in our study in the frontal, temporal and occipital regions. Possible causes for divergence could be sample size, as well as confounding effects. However, even when controlling for age, intelligence, BMI, and depression, finding remained similar. Indeed, when evaluating spatial patterns of relationships between sleep duration and cortical thickness, we observed cross-sample consistency, suggestive that the degree of impact of sleep duration in brain structure varied across samples, but that the direction of the relation between sleep and cortical thickness was comparable across the cortex. Indeed, though phenotypic relationships were diverging, 3 out of 4 local relationships between sleep and cortical thickness were driven by additive genetic factors, suggestive of a system-level impact of sleep on brain structure and function, with modest but robust local associations.

Univariate relationships between sleep, brain structure, and behavior in two independent samples were further corroborated by our multivariate analysis. We could identify multivariate, latent relationships between cortical thickness on the one hand and sleep, intelligence, BMI, and depression on the other. In the first factor, representing a positive-negative axis of behavior, where sleep duration, together with intelligence, low BMI, and high sleep quality, showed a negative relation to thickness in frontal and parietal areas, but a positive relation to sensory-motor and parahippocampal areas. Intriguingly these latent relationships were robust across samples and driven by shared additive genetic effects. Behaviorally the observed latent factor related positively to intelligence, as well as various other cognitive domains, life-satisfaction, and income and negatively to BMI, stress, smoking, and anxiety in an independent sample. Indeed, this factor mirrors the previously reported positive-negative axis of behavior previously defined in a sub-sample of the HCP using functional connectivity. Here, the axis related to increased functional connectivity of the default mode network and negative associations in sensorimotor networks ^73^. Second, we observed a second axis with a positive link between low sleep quality, high intelligence, high depression and high BMI score and thickness of dorsolateral frontal cortex, accompanied by negative relation to thickness in sensorimotor areas. This axis related again positively to intelligence and various cognitive traits, but also mental health traits such as depression, avoidance, sadness, neuroticism, and loneliness, and negatively to conscientiousness, positive affect as well as mean purpose, contrasting cognitive aptitudes such as intelligence and attention with social-emotional skills such as emotional support, friendship, and extraversion.

Our multivariate observations are broadly in line with our univariate results. Indeed, the first brain factor again highlights a negative relation between frontal thickness and sleep duration, whereas temporal-occipital regions show a positive relation with duration of sleep, reconciling at the same time divergent findings in the two independent samples. However, our latent model also provides a system-level perspective on the relation between sleep and behavior to brain structure. Here unimodal and heteromodal association cortices show an inverse relation to sleep and behavioral variability. A previous body of literature have put forward a so-called hierarchical model of brain function stretching from unimodal to transmodal cortices, enabling both externally as well as internally oriented processing ^74, 75^. Indeed, it is likely that sleep as well as intelligence, BMI, and depression do not only relate to internally oriented processes, but also functional processes focused on the external world supported by somatosensory cortices. For example, previous work has implicated sleep deprivation in sensorimotor coupling, where sleep deprived individuals showed difficulties standing upright ^76^. Next to this, memory consolidation processes during sleep has been linked to primary and secondary sensorimotor cortices. For example, in mice, inhibition of projecting axons from motor cortex to somatosensory cortex impaired sleep-dependent reactivation of sensorimotor neurons and memory consolidation ^77^. In similar vein, other studies applying multivariate methods to understand the relation between system-level brain function and complex behavior have also implicated alterations of inter-network relationships between somatosensory and heteromodal association cortices in mental function and dysfunction^78, 79^. It is possible such disruptions are due to dissociable neurodevelopmental as well as genetic effects affecting the hierarchical interrelation of these brain systems ^74^. Future research on the neurobiology of sleep therefor requires to be conducted with functional and structure connectivity data enabling more direct analysis of the relation between system-level connectivity, sleep, and behavior.

In addition to providing a novel perspective on the shared neurobiological basis of sleep, mental and physical health, we observed, in line with previous literature^17 18 43^, that variance in global sleep quality and sleep duration was in part driven by additive genetic effects. A recent GWAS study using 446,118 adults from UK Biobank identified 78 loci, mainly PAX8 locus, for self-reported habitual sleep duration^43^. Moreover, the authors observed, similar to our observations, significant genetic overlap between sleep, markers of mental and physical health, as well as education attainment. It is likely the observed genetic correlation within our sample between cortical thickness, intelligence, BMI, and sleep is due to mediated pleiotropy (a gene affects A which affects B). Thus, it could be that a genetic mechanism affects grey matter macrostructure and associated function and, as a consequence, sleep duration. Alternatively, it could be that genetic variation affects brain function which in turn modulates both macroscale structure and sleep duration, or a genetic mechanism affects sleeping behaviors through non-brain processes and in turn affects brain function and structure. However, it is worthy to mention that our genetic correlations analysis does not provide causal mechanisms on the relation between brain structure, sleep, and behavior. Indeed, there is genetic evidence for a bidirectional relationship between sleep duration and schizophrenia ^80^, as well as smoking behavior ^81^, highlighting the complex interplay between and behavior. Next to this, though there is a negative genetic correlation between short sleep duration and long sleep duration^80^, suggestive of shared biological mechanisms, it is possible both relate to partly distinct underlying biological mechanisms. Further studies will be needed to target how shorter and longer sleep differentially affects brain and behavior and could utilize longitudinal datasets with imaging and deep phenotyping to further disentangle causal relationships between sleep and brain structure and function.

Another appealing feature of our integrative perspective on sleep, behavior, and brain structure is that it may be relevant for future work targeting the biology of sleep at macroscopic, microcircuit, and molecular scales. Studies on the development have shown a close relation between sleep, behavioral problems and school performance in children ^50^. At the same time childhood is an essential time for neurodevelopment ^82, 83^ and combining these two lines of research might help to understand how sleep abnormalities and healthy sleep patterns relate to neurodevelopment in youth. At the same time, sleep disturbances have been related to neurodegenerative conditions, and may drive early-onset pathogenesis. Indeed, sleep disruption has been observed to upregulate neuronal activity upregulating the production of amyloid-beta proteins resulting in exacerbated tau pathology in various mouse models ^84^. Also, sleep disturbances in ageing can directly influences synaptic homeostasis and cognitive function. By providing novel system-level evidence integrating unimodal and heteromodal patterns in macroscale neurobiology with sleep and behavior, follow-up research could further disseminate the two brain ‘modes’ underlying sleep and general functioning during development and ageing and identify functional mechanisms that promote healthy development and ageing.

We close by noting that our findings on the interrelation between sleep, mental and physical health and brain structure was made possible by the open HCP and eNKI neuroimaging repositories. These initiatives offer cognitive neuroimaging communities an unparalleled access to large datasets for the investigation of the brain basis of individual difference. They have also enabled us to highlight variability across samples, and our work involved several statistical corrections as well as validation experiments to verify stability of our observations. Notably we observed that different samples highlight different local relationships between individual difference and brain structure, which nevertheless relate to a common neurobiological phenotype. Given that reproducibility is increasingly important nowadays, our study illustrates the advantages of open data to increase our understanding of complex traits.

## Materials and Methods

### Participants and study design: Human Connectome Project

For our analysis we used the publicly available data from the Human Connectome Project (HCP; http://www.humanconnectome.org/), which comprised data from 1206 individuals (656 females), 298 monozygotic twins (MZ), 188 dizygotic twins (DZ), and 720 singletons, with mean age 28.8 years (SD = 3.7, range = 22–37). We included for whom the scans and data had been released (humanconnectome.org) after passing the HCP quality control and assurance standards. The full set of inclusion and exclusion criteria are described elsewhere ^85^.

For our *phenotypical* analyses, we selected an unrelated subsample with complete behavioral data (n=457). After removing individuals with missing structural imaging our sample for phenotypical correlations consisted of 424 (228 females) individuals with mean age of 28.6 years (SD =3.7, range =22-36). See Table 6. For our *twin-based genetic* analyses, we used the complete sample of individuals with complete structural imaging for structural grey matter and behavioral data for sleep co-heritability analyses including 1105 individuals (599 females), 285 MZ twins, 170 DZ twins, and 650 singletons, with mean age 28.8 years (SD = 3.7, range = 22–37). Please see further Table 7. Environmental correlations were also derived in this sample as a by-product of analysis of genetic correlation.

**Table 4.** Behavioral characteristics of the HCP unrelated sample.

| Measure | n | mean±SD (range) |
| --- | --- | --- |
| *Males/Females* | 196/228 | - |
| *Age* | 424 | 28.6±3.7 (22-36) |
| *Sleep duration (hours)* | 424 | 6.8±1.2 (2.5-10) |
| *Total sleep quality* | 424 | 4.9±2.8 (0-15) |
| *BMI* | 424 | 26.6±5.3 (16.7-44.5) |
| *Intelligence (Total Cognitive Score)* | 418 | 121.5±14.7 (84.6-153.4) |
| *Depression (DSM-scale)* | 419 | 54.1±6.1 (50-87) |

**Table 5.** Behavioral characteristics of the complete HCP sample including twins and siblings.

| Measure | n | mean±SD (range) |
| --- | --- | --- |
| *Males/Females* | 507/606 | - |
| *Age* | 1113 | 28.8±3.7 (22-37) |
| *Sleep duration (hours)* | 1113 | 6.8±1.1 (2.5-12) |
| *Total sleep quality* | 1113 | 4.8±2.8 (0-19) |
| *BMI* | 1112 | 26.5±5.2 (16.5-47.8) |
| *Intelligence (Total Cognitive Score)* | 1096 | 121.8±14.6 (84.6-153.4) |
| *Depression (DSM scale)* | 1105 | 53.9±5.7 (50-87) |

**Table 6.** behavioral characteristics of the eNKI sample.

| **Measure** | **n** | **mean±SD (range)** |
| --- | --- | --- |
| *Males/Females* | 296/487 | - |
| *Age* | 783 | 41.2±20.3 (12-85) |
| *Sleep duration (hours)* | 783 | 6.9±1.3 (3-12) |
| *Total sleep quality* | 783 | 4.6±3.2 (0-17) |
| *BMI* | 757 | 27.1±5.9 (15.7-50.0) |
| *Intelligence (WASI)* | 783 | 101.9±13.3 (65-141) |
| *Depression (BDI)* | 468 | 7.0±6.9 (1-40) |

### Structural imaging processing: Human Connectome Project

MRI protocols of the HCP are previously described^86, 87^. In particular, the applied pipeline to obtain the Freesurfer-segmentation is described earlier^86^ and is recommended for the HCP data. The pre-processing steps included co-registration of T1 and T2 images, B1 (bias field) correction, and segmentation and surface reconstruction using FreeSurfer version 5.3-HCP to estimate cortical thickness ^86^.

### Participants and study design: eNKI sample

To evaluate the cross-sample reproducibility of observations we additionally investigated correspondence between sleep and cortical brain structure in the enhanced Nathan Kline Institute-Rockland Sample (NKI). The sample was made available by the Nathan-Kline Institute (NKY, NY, USA), as part of the ‘*enhanced NKI-Rockland sample*’ (https://www.ncbi.nlm.nih.gov/pmc/articles/PMC3472598/). All approvals regarding human subjects’ studies were sought following NKI procedures. Scans were acquired from the International Neuroimaging Data Sharing Initiative (INDI) online database http://fcon_1000.projects.nitrc.org/indi/enhanced/studies.html

For our *phenotypical* analyses, we selected individuals with complete sleep and imaging data. Our sample for phenotypical correlations consisted of 783 (487 females) individuals with mean age of 41.2 years (SD =20.3, range =12-85). Please see Table 8 for demographic characteristics.

### Structural imaging processing: NKI Rockland sample

3D magnetization-prepared rapid gradient-echo imaging (3D MP-RAGE) structural scans^88^ were acquired using a 3.0 T Siemens Trio scanner with TR=2500 ms, TE=3.5 ms, Bandwidth=190 Hz/Px, field of view=256 × 256 mm, flip angle=8°, voxel size=1.0 × 1.0 × 1.0 mm. More details on image acquisition are available at http://fcon_1000.projects.nitrc.org/indi/enhanced/studies.html. All T1 scans were visually inspected to ensure the absence of gross artefacts and subsequently pre-processed using the Freesurfer software library (http://surfer.nmr.mgh.harvard.edu/) Version 5.3.0^89^.

### Parcellation-summaries of cortical thickness

We used a parcellation scheme^47^ based on the combination of a local gradient approach and a global similarity approach using a gradient-weighted Markov Random models. The parcellation has been extensively evaluated with regards to stability and convergence with histological mapping and alternative parcellations. In the context of the current study, we focus on the granularity of 200 parcels. In order to improve signal-to-noise and improve analysis speed, we opted to average unsmoothed structural data within each parcel. Thus, cortical thickness of each region of interest (ROI) was estimated as the trimmed mean (10 percent trim).

### Selection of behavioral markers based on HCP phenotypic traits

First, to constrain analyses we selected primary markers for cognition, mental and physical health based on the relation of sleep to these traits in HCP. The selected traits include 38 emotional, cognitive, NEO-FFI personality, as well as the 7 PSQI sleep markers for reference, based on the unrestricted phenotypic data as well as 46 mental and physical health markers based on the restricted phenotypic data. For more information on available phenotypes, please see: https://wiki.humanconnectome.org/display/PublicData/HCP+Data+Dictionary+Public-+Updated+for+the+1200+Subject+Release#HCPDataDictionaryPublic-Updatedforthe1200SubjectRelease-Instrument:SocialRelationships.

### Behavioral markers: HCP

Inter-individual difference in sleep quality were derived from information of the self-reported Pittsburg Sleep Questionnaire (PSQI)^11^, which is common measure of sleep quality with significant item-level reliability and validity. For markers of life function, we used BMI (703 * weight/ (height)^2) and the ASR depression DSM-oriented scale for Ages 18-59^46^ (https://aseba.org/). The ASR is a self-administered test examining diverse aspects of adaptive functioning and problems. Scales are based on 2020 referred adults and normed on 1767 non-referred adults. The test-retest reliability of the ASR was supported by 1-week test-retest that were all above 0.71. The ASR also has good internal consistency (0.83), in the current study we focused on depression sub-score. As a proxy for intelligence we used the NIH Toolbox Cognition^44^, ‘total composite score’. The Cognitive Function Composite score is derived by averaging the normalized scores of each of the Fluid and Crystallized cognition measures, then deriving scale scores based on this new distribution. Higher scores indicate higher levels of cognitive functioning. Participant score is normed to those in the entire NIH Toolbox Normative Sample (18 and older), regardless of age or any other variable, where a score of 100 indicates performance that was at the national average and a score of 115 or 85, indicates performance 1 SD above or below the national average.

### Behavioral markers: NKI

Sleep markers were derived from the Pittsburg Sleep Questionnaire (see further the section on this question in the HCP sample). Intelligence was measured using the Wechsler Abbreviated Scale of Intelligence (WASI-II). The WASI is a general intelligence, or IQ test designed to assess specific and overall cognitive capabilities and is individually administered to children, adolescents and adults (ages 6-89). It is a battery of four subtests: Vocabulary (31-item), Block Design (13-item), Similarities (24-item) and Matrix Reasoning (30-item). In addition to assessing general, or Full Scale, intelligence, the WASI is also designed to provide estimates of Verbal and Performance intelligence consistent with other Wechsler tests. Specifically, the four subtests comprise the full scale and yield the Full Scale IQ (FSIQ-4). The Vocabulary and Similarities subtests are combined to form the Verbal Scale and yield a Verbal IQ (VIQ) score, and the Block Design and Matrix Reasoning subtests form the Performance Scale and yield a Performance IQ (PIQ) score ^45^.

Depression was measured using the Beck Depression Inventory (BDI – II). The BDI-II is a 21-item self-report questionnaire assessing the current severity of depression symptoms in adolescents and adults (ages 13 and up). It is not designed to serve as an instrument of diagnosis, but rather to identify the presence and severity of symptoms consistent with the criteria of the DSM-IV. Questions assess the typical symptoms of depression such as mood, pessimism, sense of failure, self-dissatisfaction, guilt, punishment, self-dislike, self-accusation, suicidal ideas, crying, irritability, social withdrawal, insomnia, fatigue, appetite, and loss of libido. Participants are asked to pick a statement on a 4-point scale that best describes the way they have been feeling during the past two weeks ^90^.

Body-mass-index was calculated using weight and height. These vitals are obtained and recorded by study staff. Height was recorded in centimeters. Weight was recorded in kilograms. Body Mass Index (BMI) was automatically calculated.

### Statistical analysis

#### Phenotypical analysis

For our phenotypical analysis in the HCP sample we selected an unrelated subsample to overcome possible bias due to genetic similarity of individuals. To assess phenotypical relationships between sleep parameters and behavior/brain structure, we used Spearman’s correlation test to account for outliers, while controlling for age, sex, age × sex, age × age. In our structural whole-brain analysis, we additionally controlled for global thickness. Findings were similar when controlling for depression, BMI or intelligence additionally. We controlled for multiple comparisons at 5% Bonferroni, per analysis step of univariate behavior and univariate brain analysis, controlling for the number of tests at each step, and report exact FDRq thresholds for reference. We used the Robust Correlation Toolbox for Matlab to define confidence intervals in our post-hoc phenotypic correlations (Pernet, 2013).

#### Heritability and genetic correlation analysis

To investigate the heritability and genetic correlation of sleep parameters, and brain structure, we analyzed 200 parcels of cortical thickness, as well as sleep parameters of each subject in a twin-based heritability analysis. As previously described^91^, the quantitative genetic analyses were conducted using Sequential Oligogenic Linkage Analysis Routines (SOLAR)^92^. SOLAR uses maximum likelihood variance-decomposition methods to determine the relative importance of familial and environmental influences on a phenotype by modeling the covariance among family members as a function of genetic proximity. This approach can handle pedigrees of arbitrary size and complexity and thus, is optimally efficient with regard to extracting maximal genetic information. To ensure that neuroimaging traits, parcels of cortical thickness, conform to the assumptions of normality, an inverse normal transformation was applied^91^.

Heritability (*h_2_*) represents the portion of the phenotypic variance (σ_2_p) accounted for by the total additive genetic variance (σ_2_g), i.e., *h_2_* = σ_2_g/σ_2_p. Phenotypes exhibiting stronger covariances between genetically more similar individuals than between genetically less similar individuals have higher heritability. Within SOLAR, this is assessed by contrasting the observed covariance matrices for a neuroimaging measure with the structure of the covariance matrix predicted by kinship. Heritability analyses were conducted with simultaneous estimation for the effects of potential covariates. For this study, we included covariates including age, sex, age × sex interaction, age2, age2 × sex interaction. When investigating cortical thickness, we additionally controlled for global thickness effects as well as depression score, BMI, and intelligence in post-hoc tests. Heritability estimates were corrected for multiple comparisons at 5% Bonferroni, controlling for the number of parcels.

To determine if variations in sleep and brain structure were influenced by the same genetic factors, genetic correlation analyses were conducted. Specifically, bivariate polygenic analyses were performed to estimate genetic (ρ_g_) and environmental (ρ_e_) correlations, based on the phenotypical correlation (ρ_p_), between brain structure and sleep with the following formula: ρ_p_ = ρ_g_√(*h*_21_*h*_22_) + ρ_e_√[(1 − *h*_21_)(1 − *h*_22_)], where *h*_21_ and *h*_22_ are the heritability’s of the parcel-based cortical thickness and the sleep parameters. The significance of these correlations was tested by comparing the log likelihood for two restricted models (with either ρ_g_ or ρ_e_ constrained to be equal to 0) against the log likelihood for the model in which these parameters were estimated. A significant genetic correlation (using a 5% Bonferroni) is evidence suggesting that both phenotypes are influenced by a gene or set of genes^93^.

### Partial least squares

PLS is a multivariate data-driven statistical technique that aims to maximize the covariance between two matrices by deriving *latent components* (LCs), which are optimal linear combinations of the original matrices ^94, 95^. We applied PLS to the cortical thickness and personality measures of all participants. In short, PLS performs data normalization, cross-covariance, and singular value decomposition. Following, brain and behavioral scores are created and premutation testing is performed to assess significance of each latent factor solution. Last, bootstrapping is performed to test the stability of the brain saliencies.

Each LC is characterized by a distinct cortical thickness pattern (called brain *saliences*) and a distinct behavioral profile (called behavioral *saliences*). By linearly projecting the brain and behavioral measures of each participant onto their respective saliences, we obtain individual-specific brain and behavioral *composite scores* for each LC. By construction, PLS seeks to find saliences that maximize across-participant covariance between the brain and behavioral composite scores. The number of significant LCs was determined by a permutation (1000 permutations). The p-values (from the permutation test) for the LCs were corrected for multiple comparisons using a false discovery rate (FDR) of p < 0.05. To interpret the LCs, we further computed Pearson’s correlations between behavioral composite scores for each LC and behavioral measures. A large positive (or negative) correlation for a particular behavioral measure for a given LC indicates greater importance of the behavioral measure for the LC ^79^. For the brain saliencies, though all regions contributed to the latent brain score, we highlighted regions with a bootstrap ratio > 2, approximately p<0.05. Findings where summarized at the level of macroscale function networks^49^, by averaging the BSR score per network, as well as summarizing the relative contribution of each functional network to positive (BSR>2) as well as negative (BSR<-2) relations. Here we controlled for the size of the network.

### Functional decoding

All significant parcels were functionally characterized using the Behavioral Domain meta-data from the BrainMap database using forward inference (http://www.brainmap.org)^96, 97^. To do so, volumetric counterparts of the surface-based parcels were identified. In particular, we identified those meta-data labels (describing the computed contrast [behavioral domain]) that were significantly more likely than chance to result in activation of a given parcel^98–100^. That is, functions were attributed to the identified effects by quantitatively determining which types of experiments are associated with activation in the respective parcellation region. Significance was established using a binomial test (p < 0.05, corrected for multiple comparisons using false discovery rate (FDR)).

## SUPPLEMENT

### Supplementary Figures

**Figure S1.**
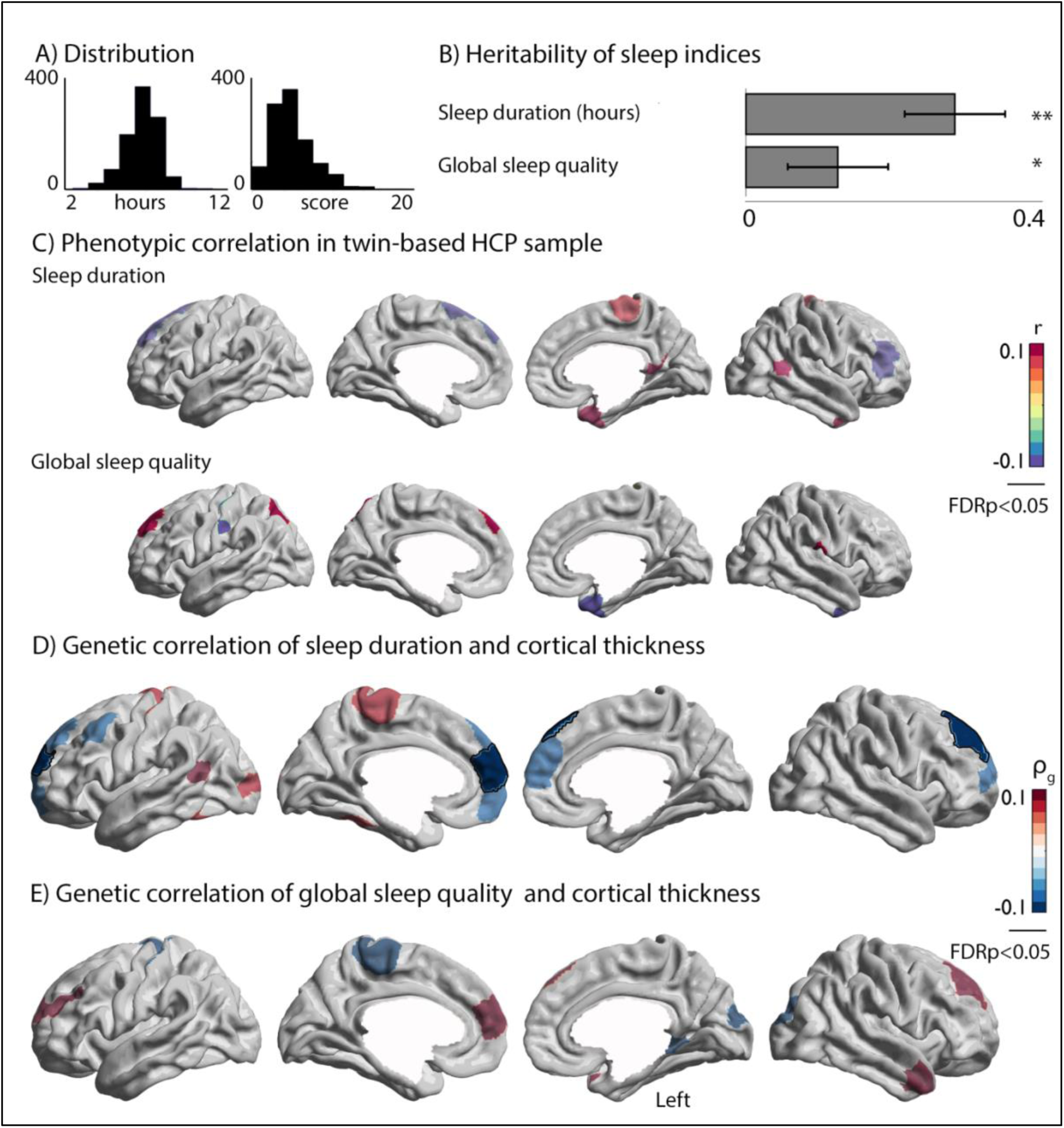
Shared genetic basis of sleep duration and cortical thickness. A) Distribution of variables; B) Heritability of sleep duration and global sleep quality score. Significant (p<0.05/2) heritability is annotated with **, and p<0.05 with *; C). Phenotypic correlation of sleep duration and global sleep quality in the related sample; D+E) genetic correlation of sleep duration/global sleep quality and voxel-wise cortical thickness. Significant associations between sleep indices and brain structure have black outline, whereas trends (p<0.01) are transparent. Red indicates a positive relationship, whereas blue indicates a negative genetic relationship between sleep and brain structure.

**Figure S2.**
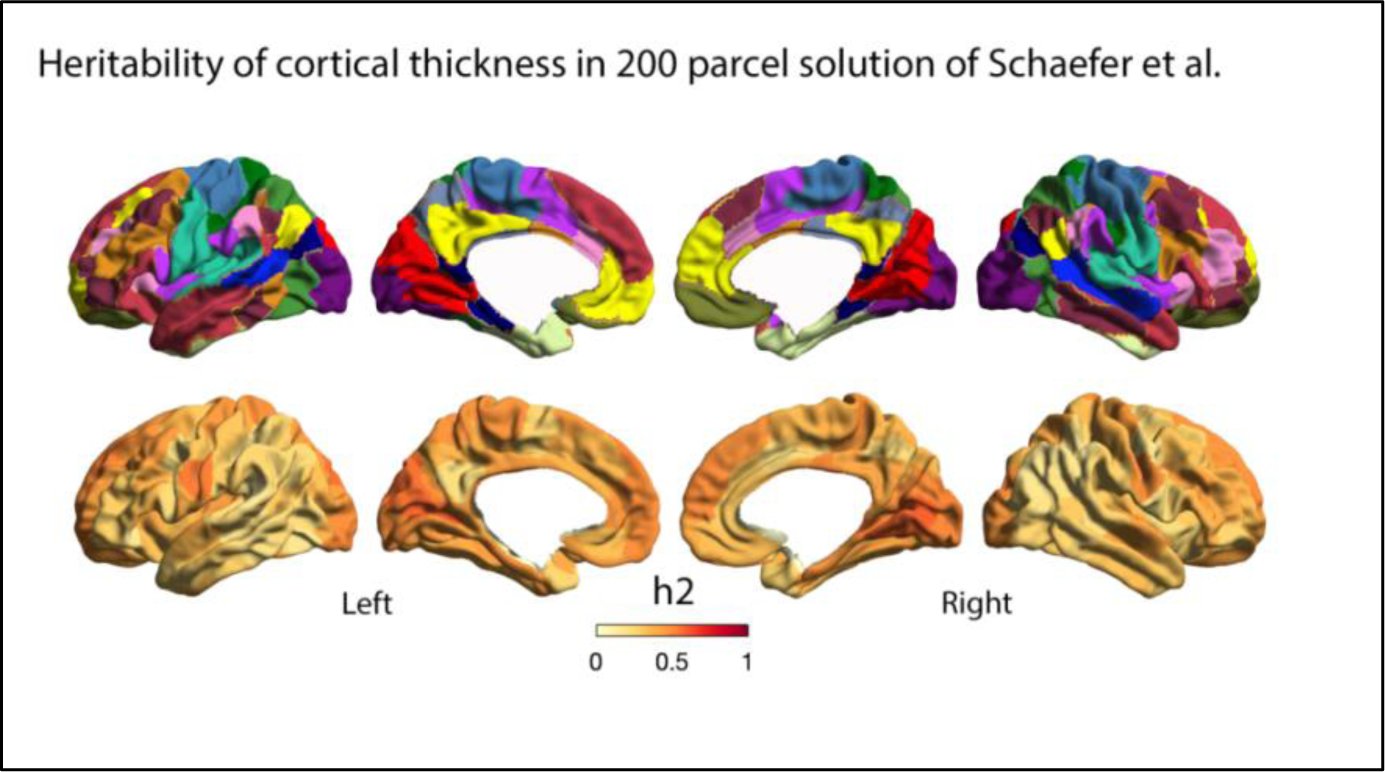
Heritability of cortical thickness. Heritability of brain structure (lower panel) in the 200-parcel solution of Schaefer atlas^34^ (upper panel). The 200-parcel solution by Schaefer et al is based on both local and global functional connectivity. In each parcel, we extracted trimmed mean (10%) cortical thickness values of each individual. Subsequently, we computed heritability of variance in cortical thickness in each region and displayed this on the cortex. Please see Supplementary Table 7 for heritability values as well as standard errors and significance.

**Figure S3.**
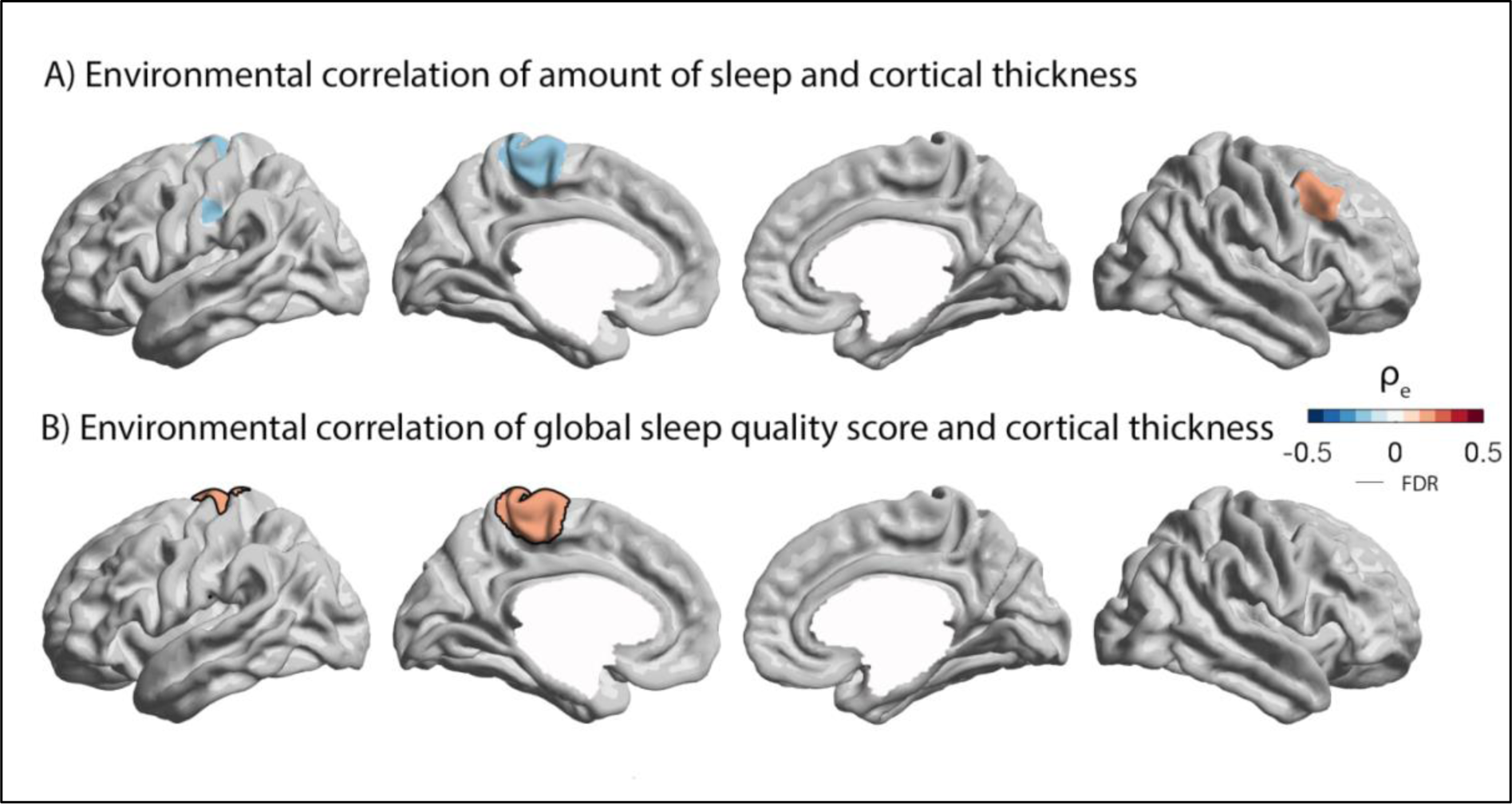
Environmental relationship between sleep and brain structure. A) Environmental correlation between sleep duration and brain structure and B) global sleep quality and brain structure. Significant associations between sleep indices and brain structure have black outline, whereas trends (p<0.01) are transparent. Red indicates a positive relationship, whereas blue indicates a negative genetic relationship between sleep and brain structure.

**Figure S4.**
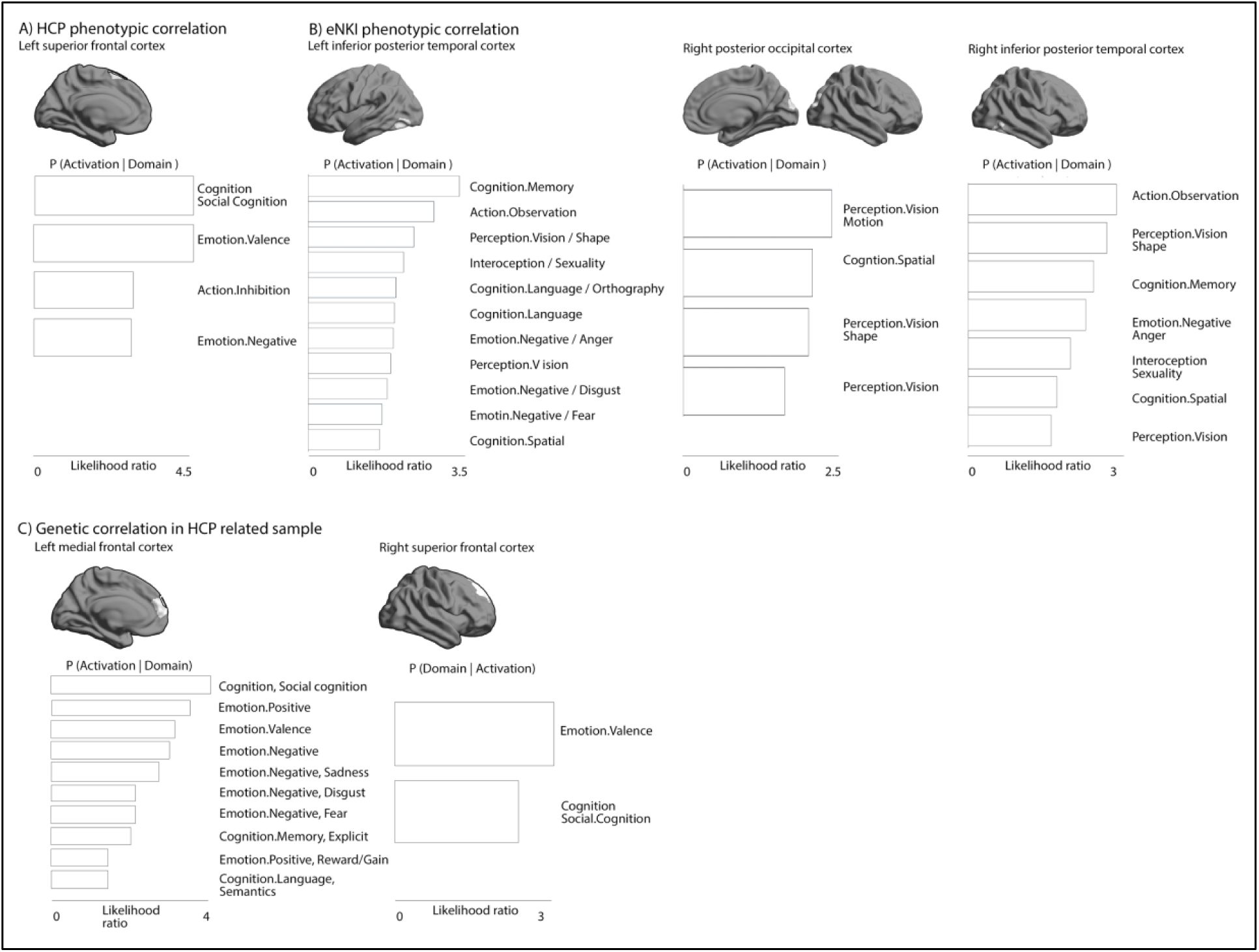
Cortical thickness correlates of sleep duration are functionally linked with cognitive and emotion processing. Quantitative functional decomposition of brain markers of sleep based on the FDR-corrected clusters of phenotypical/genetic relationships shown in Fig 1 and Fig S1.

### Supplementary Tables

**Table S1.**
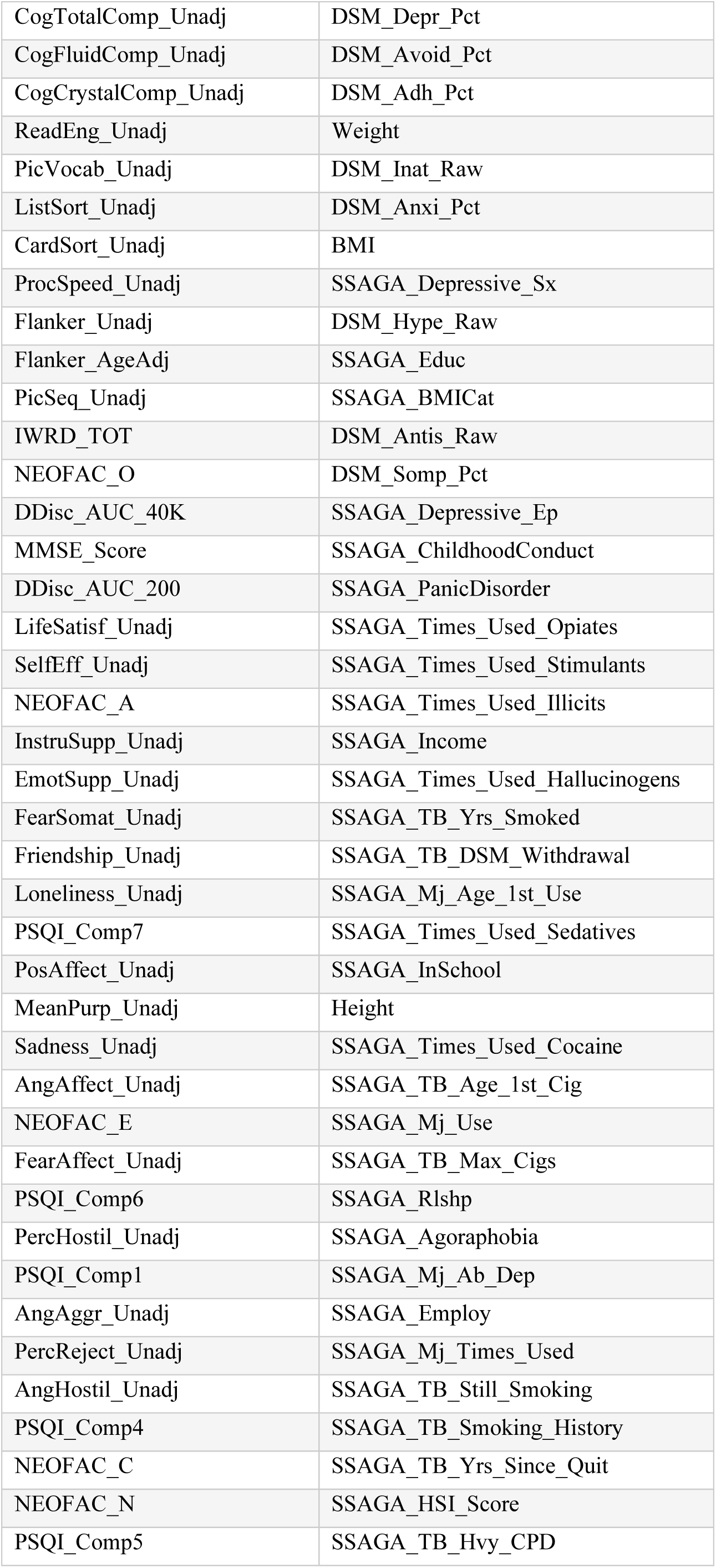
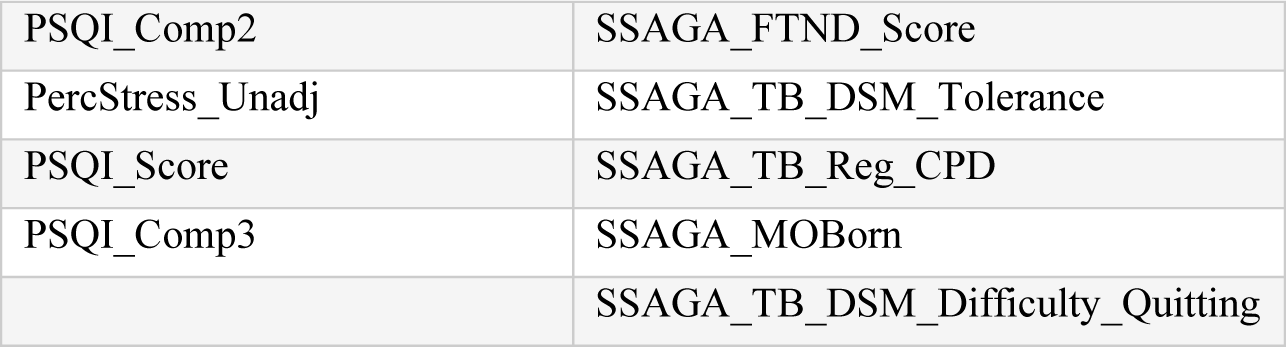
List of selected behavior markers in HCP sample. Behavioral markers to evaluate the relation between sleep and cognition, mental, and physical health. For further details, please see: https://wiki.humanconnectome.org/display/PublicData/HCP+Data+Dictionary+Public-+Updated+for+the+1200+Subject+Release#HCPDataDictionaryPublic-Updatedforthe1200SubjectRelease-Instrument:SocialRelationships

|  |  |
| --- | --- |
| CogTotalComp_Unadj | DSM_Depr_Pct |
| CogFluidComp_Unadj | DSM_Avoid_Pct |
| CogCrystalComp_Unadj | DSM_Adh_Pct |
| ReadEng_Unadj | Weight |
| PicVocab_Unadj | DSM_Inat_Raw |
| ListSort_Unadj | DSM_Anxi_Pct |
| CardSort_Unadj | BMI |
| ProcSpeed_Unadj | SSAGA_Depressive_Sx |
| Flanker_Unadj | DSM_Hype_Raw |
| Flanker_AgeAdj | SSAGA_Educ |
| PicSeq_Unadj | SSAGA_BMICat |
| IWRD_TOT | DSM_Antis_Raw |
| NEOFAC_O | DSM_Somp_Pct |
| DDisc_AUC_40K | SSAGA_Depressive_Ep |
| MMSE_Score | SSAGA_ChildhoodConduct |
| DDisc_AUC_200 | SSAGA_PanicDisorder |
| LifeSatisf_Unadj | SSAGA_Times_Used_Opiates |
| SelfEff_Unadj | SSAGA_Times_Used_Stimulants |
| NEOFAC_A | SSAGA_Times_Used_Illicits |
| InstruSupp_Unadj | SSAGA_Income |
| EmotSupp_Unadj | SSAGA_Times_Used_Hallucinogens |
| FearSomat_Unadj | SSAGA_TB_Yrs_Smoked |
| Friendship_Unadj | SSAGA_TB_DSM_Withdrawal |
| Loneliness_Unadj | SSAGA_Mj_Age_1st_Use |
| PSQI_Comp7 | SSAGA_Times_Used_Sedatives |
| PosAffect_Unadj | SSAGA_InSchool |
| MeanPurp_Unadj | Height |
| Sadness_Unadj | SSAGA_Times_Used_Cocaine |
| AngAffect_Unadj | SSAGA_TB_Age_1st_Cig |
| NEOFAC_E | SSAGA_Mj_Use |
| FearAffect_Unadj | SSAGA_TB_Max_Cigs |
| PSQI_Comp6 | SSAGA_Rlshp |
| PercHostil_Unadj | SSAGA_Agoraphobia |
| PSQI_Comp1 | SSAGA_Mj_Ab_Dep |
| AngAggr_Unadj | SSAGA_Employ |
| PercReject_Unadj | SSAGA_Mj_Times_Used |
| AngHostil_Unadj | SSAGA_TB_Still_Smoking |
| PSQI_Comp4 | SSAGA_TB_Smoking_History |
| NEOFAC_C | SSAGA_TB_Yrs_Since_Quit |
| NEOFAC_N | SSAGA_HSI_Score |
| PSQI_Comp5 | SSAGA_TB_Hvy_CPD |
| PSQI_Comp2 | SSAGA_FTND_Score |
| PercStress_Unadj | SSAGA_TB_DSM_Tolerance |
| PSQI_Score | SSAGA_TB_Reg_CPD |
| PSQI_Comp3 | SSAGA_MOBorn |
|  | SSAGA_TB_DSM_Difficulty_Quitting |

**Table S2.** Phenotypic association between sleep duration and behavior in HCP sample. Speaman’s rho between sleep duration and the respective behavioral score in HCP, controlled for the number of tests.

|  |  |
| --- | --- |
| SSAGA_Educ | r=0.13 |
| **BMI** | r=-0.13 |
| DSM_Antis_Raw | r=-0.14 |
| NEOFAC_A | r=0.20 |
| DDisc_AUC_40K | r=0.19 |
| DDisc_AUC_200 | r=0.16 |
| LifeSatisf_Unadj | r=0.14 |
| ReadEng_Unadj | r=0.13 |
| CogCrystalComp_Unadj | r=0.13 |
| PicVocab_Unadj | r=0.11 |
| **CogTotalComp_Unadj** | r=0.10 |
| AngAffect_Unadj | r=-0.11 |
| PSQI_Comp2 | r=-0.11 |
| PSQI_Comp5 | r=-0.12 |
| PercStress_Unadj | r=-0.12 |
| PSQI_Comp7 | r=-0.14 |
| PSQI_Comp1 | r=-0.36 |
| PSQI_Comp4 | r=-0.39 |
| PSQI_Score | r=-0.55 |
| PSQI_Comp3 | r=-0.89 |

**Table S3.** Phenotypic association between global sleep quality and behavior in HCP sample. Speaman’s rho between sleep quality and the respective behavioral score in HCP, controlled for the number of tests.

|  |  |
| --- | --- |
| PSQI_Score | r=1.00 |
| PSQI_Comp1 | r=0.67 |
| PSQI_Comp3 | r=0.58 |
| PSQI_Comp2 | r=0.58 |
| PSQI_Comp4 | r=0.57 |
| PSQI_Comp5 | r=0.49 |
| PSQI_Comp7 | r=0.45 |
| PSQI_Comp6 | r=0.35 |
| PercStress_Unadj | r=0.29 |
| AngAffect_Unadj | r=0.29 |
| NEOFAC_N | r=0.27 |
| FearAffect_Unadj | r=0.24 |
| AngHostil_Unadj | r=0.22 |
| Sadness_Unadj | r=0.22 |
| FearSomat_Unadj | r=0.21 |
| PercReject_Unadj | r=0.20 |
| Loneliness_Unadj | r=0.19 |
| AngAggr_Unadj | r=0.16 |
| PercHostil_Unadj | r=0.15 |
| **CogTotalComp_Unadj** | *r=-0.10* |
| PicVocab_Unadj | r=-0.12 |
| NEOFAC_E | r=-0.12 |
| ReadEng_Unadj | r=-0.12 |
| CogCrystalComp_Unadj | r=-0.13 |
| NEOFAC_C | r=-0.13 |
| Friendship_Unadj | r=-0.14 |
| EmotSupp_Unadj | r=-0.16 |
| DDisc_AUC_40K | r=-0.17 |
| DDisc_AUC_200 | r=-0.17 |
| MeanPurp_Unadj | r=-0.18 |
| PosAffect_Unadj | r=-0.20 |
| NEOFAC_A | r=-0.22 |
| LifeSatisf_Unadj | r=-0.25 |
| **DSM_Depr_Pct** | r=0.35 |
| DSM_Anxi_Pct | r=0.30 |
| DSM_Hype_Raw | r=0.24 |
| DSM_Adh_Pct | r=0.23 |
| DSM_Antis_Raw | r=0.23 |
| DSM_Somp_Pct | r=0.23 |
| DSM_Inat_Raw | r=0.20 |
| SSAGA_Depressive_Sx | r=0.17 |
| SSAGA_ChildhoodConduct | r=0.14 |
| SSAGA_Depressive_Ep | r=0.14 |
| SSAGA_TB_Smoking_History | r=0.13 |
| SSAGA_Mj_Use | r=0.13 |
| SSAGA_Mj_Times_Used | r=0.12 |
| DSM_Avoid_Pct | r=0.12 |
| SSAGA_TB_Still_Smoking | r=0.12 |
| SSAGA_BMICat | r=0.11 |
| **BMI** | r=0.11 |
| SSAGA_Income | r=-0.12 |
| SSAGA_Educ | r=-0.14 |

**Table S4.** Phenotypic and genetic correlations between confounding effects on sleep (Depression, BMI, and IQ). **FDRq<0.05, and *p<0.05

| Interrelations HCP all |  |  |  |
| --- | --- | --- | --- |
|  | Depression | BMI | IQ |
| *Depression* | 1 |  |  |
| *BMI* | 0.05 | 1 |  |
| *IQ* | -0.02 | -0.17 ** | 1 |
| Interrelations HCP unrelated subsample |  |  |  |
|  | Depression | BMI | IQ |
| *Depression* | 1 |  |  |
| *BMI* | 0.21 ** | 1 |  |
| *IQ* | 0.00 | -0.19 ** | 1 |
| Interrelations: eNKI |  |  |  |
|  | Depression | BMI | IQ |
| *Depression* | 1 |  |  |
| *BMI* | 0.12 ** | 1 |  |
| *IQ* | -0.11 ** | -0.15 ** | 1 |
| Interrelations: genetic correlation (HCP) |  |  |  |
|  | Depression | BMI | IQ |
| *Depression* | 1 |  |  |
| *BMI* | -0.02 [0.12] | 1 |  |
| *IQ* | 0.02 [0.10] | -0.27 [0.06] ** | 1 |
| Interrelations: environmental correlation (HCP) |  |  |  |
|  | Depression | BMI | IQ |
| *Depression* | 1 |  |  |
| *BMI* | 0.12 [0.07] | 1 |  |
| *IQ* | -0.08 [0.24] | 0.10 [0.07] | 1 |

**Table S5.** Intercorrelations of spatial correspondences between sleep and cortical thickness. Cross-sample replication of spatial distribution from phenotypic and genetic correlational analysis in F1 and FS1.* indicates p<0.05, ** indicates FDRq<0.05.

| **Sleep duration** | HCP (unrelated sample) | HCP (unrelated sample) | HCP (unrelated sample) | HCP (unrelated sample) |
| --- | --- | --- | --- | --- |
| HCP (unrelated sample) | 1 |  |  |  |
| HCP (total sample) | 0.77 [0.69 0.83] ** | 1 |  |  |
| *eNKI* | 0.30 [0.16 0.44] ** | 0.16 [0.02 0.29] ** | 1 |  |
| Genetic | 0.53 [0.42 0.63] ** | 0.61 [0.51 0.69] ** | 0.24 [0.12 0.37] ** | 1 |
| **Global sleep quality** | **HCP (unrelated sample)** | **HCP (unrelated sample)** | **HCP (unrelated sample)** | **HCP (unrelated sample)** |
| HCP (unrelated sample) | 1 |  |  |  |
| HCP (total sample) | 0.64 [0.54 0.72] ** | 1 |  |  |
| eNKI | 0.13 [-0.01 0.26] | -0.09 [-0.22 0.06] | 1 |  |
| Genetic | 0.19. [0.05. 0.32] ** | 0.48 [0.36 0.59] ** | -0.02 [-0.15 0.12] | 1 |

**Table S6.** Intelligence and sleep duration both have phenotypic and genetic correlation with frontal regions. Cross-sample analysis of genetic correlational analysis between frontal regions genetically associated with sleep duration on the one hand, and depression, BMI, and IQ on the other. ** reflects FDRq<0.05, * reflects p<0.05.

| Phenotypical correlation between sleep duration and cortical thickness (Fig. 1A, HCP unrelated) |  |  |  |
| --- | --- | --- | --- |
| Left superior frontal gyrus h2 = 0.46 (0.05) |  |  |  |
| Sample | Depression | BMI | IQ |
| HCP (unrelated sample) | 0.02 [-0.07 0.12] | 0.09 [-0.01 0.17] | -0.04 [-0.15 0.05] |
| eNKI | -0.04 [-0.13 0.05] | -0.01 [-0.09 0.05] | -0.08 [-0.15 -0.01] * |
| HCP (total sample) | 0.05 [-0.00 0.11] | 0.04 [-0.01 0.10] | -0.11 [-0.17 -0.05] ** |
| Genetic | -0.00 (0.13) | 0.05 (0.08) | 0.19 (0.08) * |
| Enviorment | 0.11 (0.06) | 0.04 (0.07) | 0.04 (0.07) |
| Phenotypical correlation between sleep duration and cortical thickness, (Fig. 1B, eNKI) |  |  |  |
| Left inferior temporal cortex h2 = 0.17 (0.06) |  |  |  |
| Sample | Depression | BMI | IQ |
| HCP (unrelated sample) | -0.08 [-0.18 0.01] | 0.02 [-0.07 0.12] | -0.01 [-0.11 0.09] |
| eNKI | -0.02 [-0.12 0.07] | -0.06 [-0.13 0.01] | -0.03 [-0.10 0.04] |
| HCP (total sample) | -0.07 [-0.13 0.00] * | 0.02 [-0.04 0.08] | 0.01 [-0.05 0.07] |
| Genetic | 0.001 (0.22) | 0.27 (0.14) * | -0.002 (0.12) |
| Environment | -0.07 (0.06) | -0.13 (0.07) * | 0.007 (0.06) |
| Right occipital cortex h2 = 0.51 (0.05) |  |  |  |
| Sample | Depression | BMI | IQ |
| HCP (unrelated sample) | -0.06 [-0.15 0.04] | -0.02 [-0.12 0.08] | -0.02 [-0.12 0.08] |
| eNKI | -0.03 [-0.11 0.05] | 0.07 [0.00 0.14] * | -0.02 [-0.09 0.05] |
| HCP (total sample) | -0.03 [-0.08 0.03] | -0.00 [-0.06 0.06] | 0.06 [-0.01 0.12] |
| Genetic | -0.03 (0.13) | 0.01 (0.07) | 0.19 (0.07) * |
| Environment | -0.04 (0.06) | -0.02 (0.07) | -0.15 (0.07) * |
| Right inferior temporal cortex h2 = 0.29 (0.06) |  |  |  |
| Sample | Depression | BMI | IQ |
| HCP (unrelated sample) | -0.05 [-0.14 0.05] | 0.08 [-0.01 0.18] | -0.01 [-0.10 0.08] |
| eNKI | -0.02 [-0.11 0.07] | -0.08 [-0.14 -0.01] * | 0.12 [0.05 0.19] ** |
| HCP (total sample) | -0.01 [-0.07 0.05] | 0.01 [-0.05 0.07] | 0.02 [-0.04 0.08] |
| Genetic | 0.08 (0.17) | 0.10 (0.10) | 0.20 (0.09) * |
| Environment | -0.02 (0.06) | -0.08 (0.07) | -0.07 (0.06) |

**Table S7.**
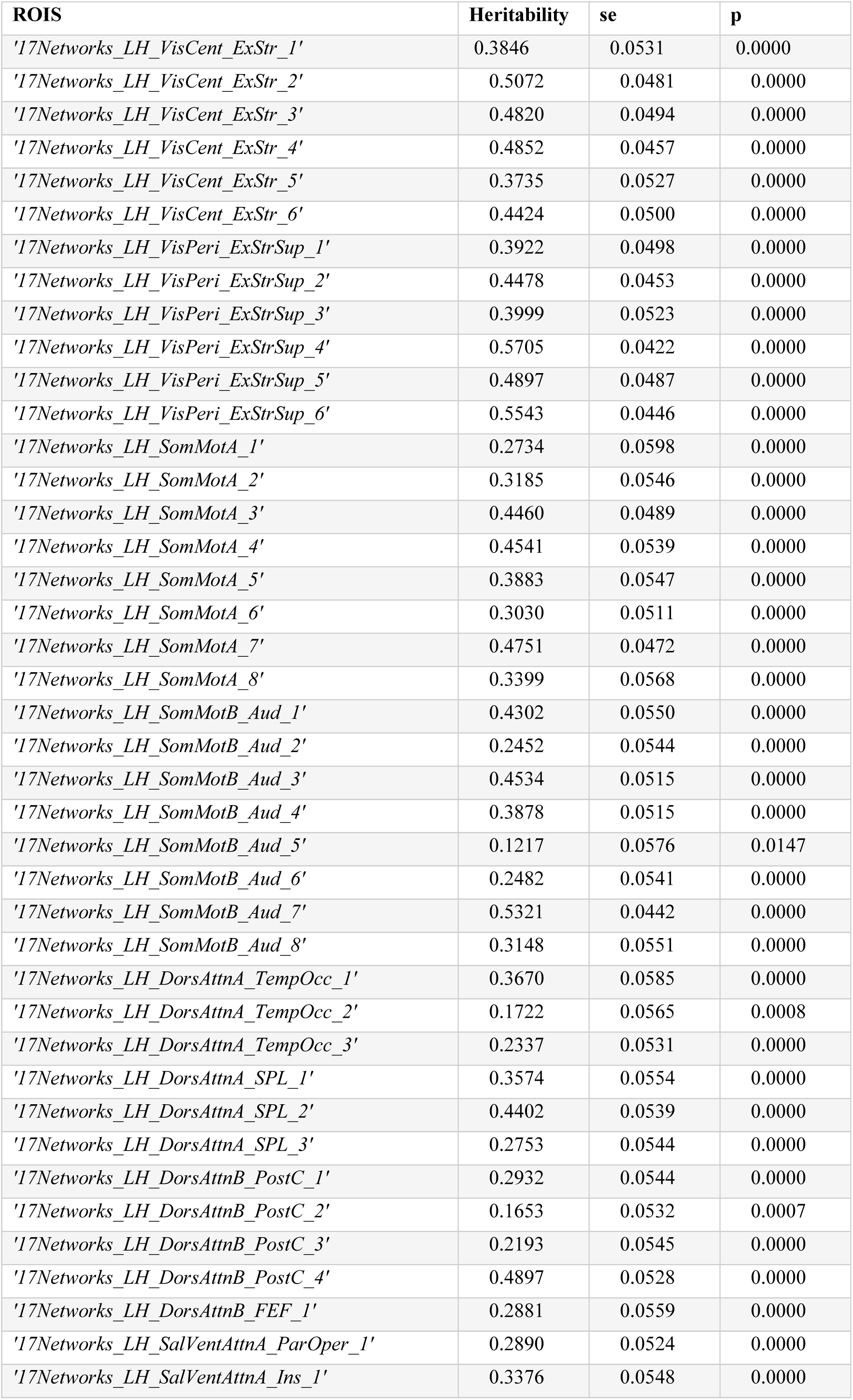
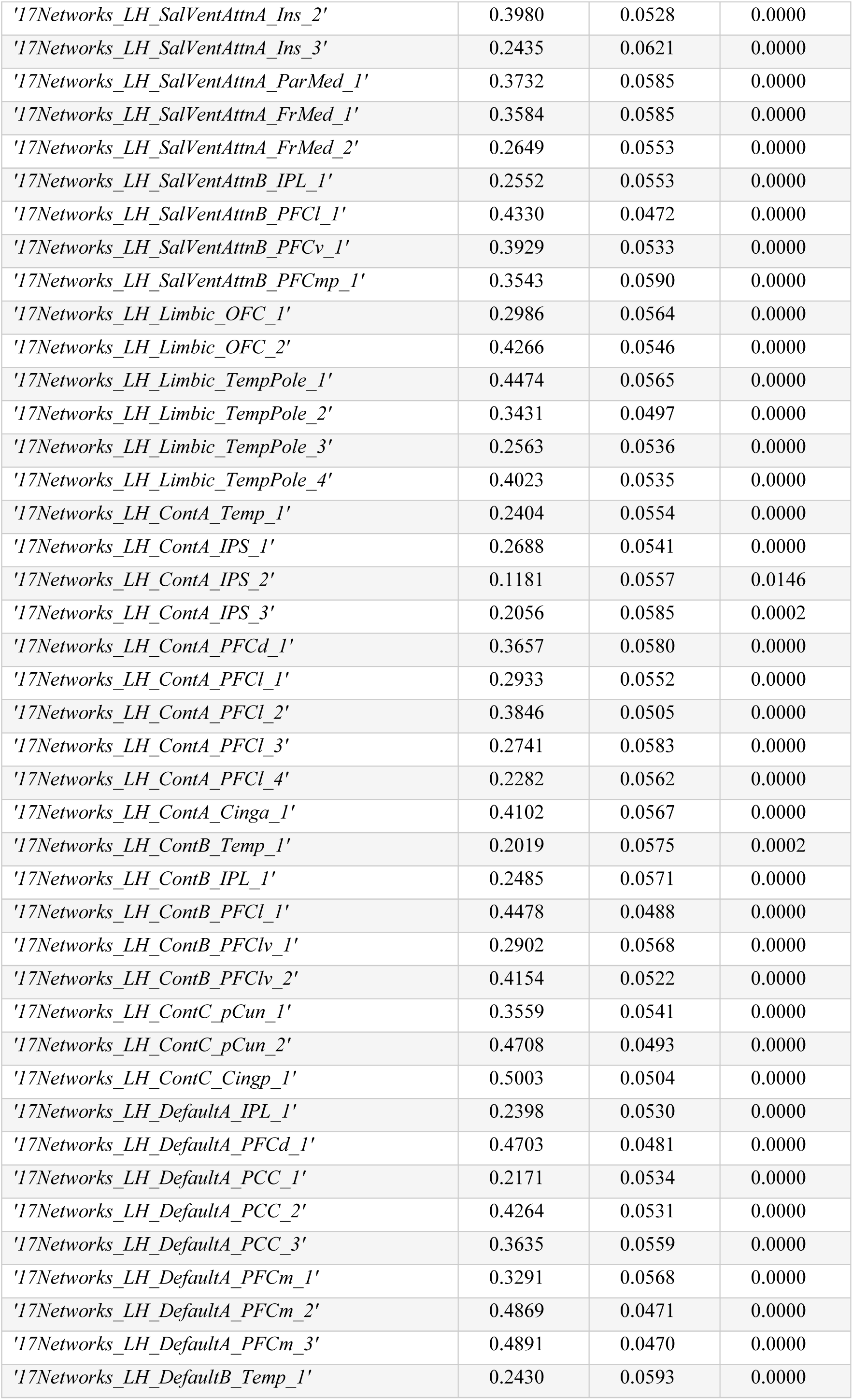
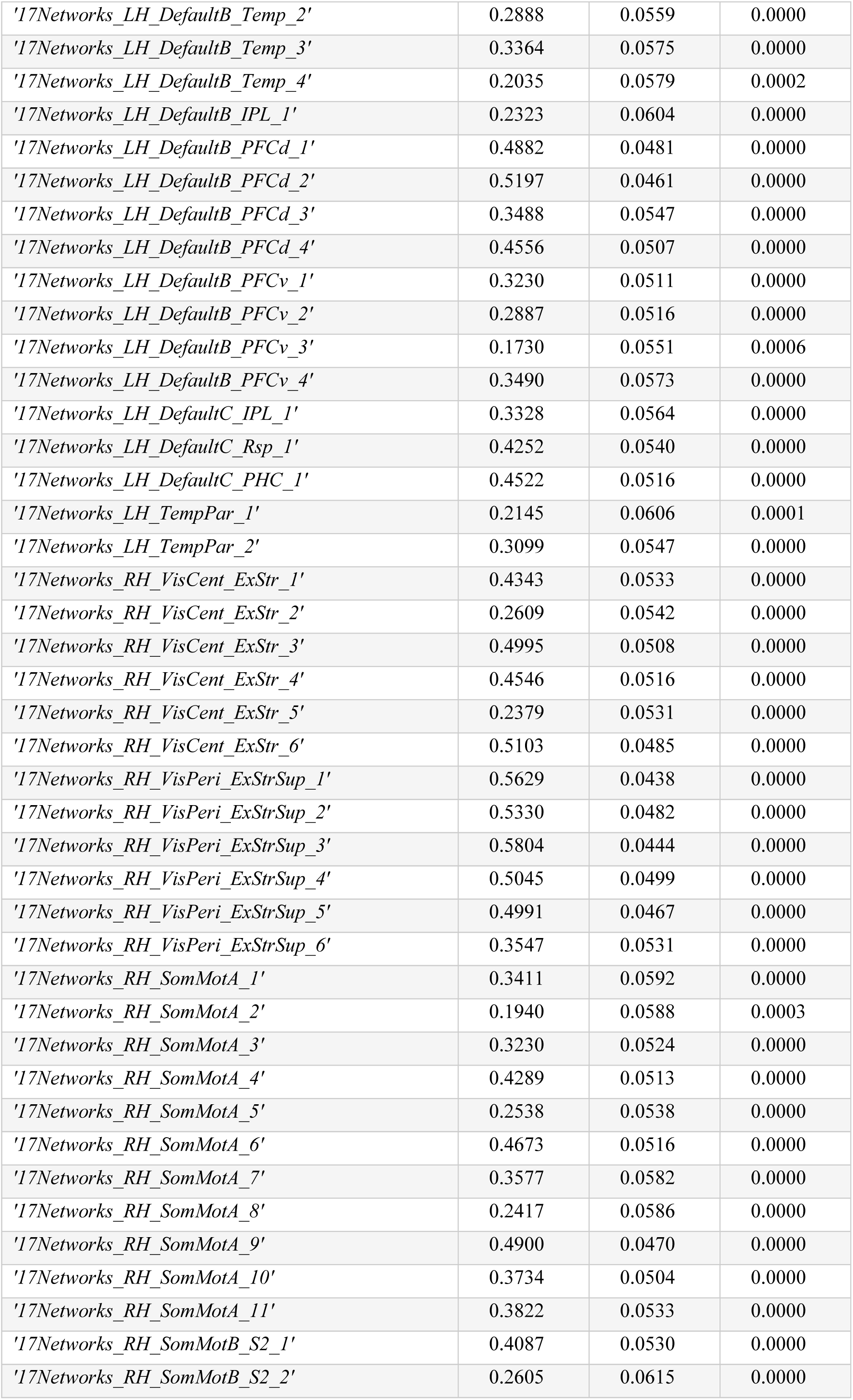
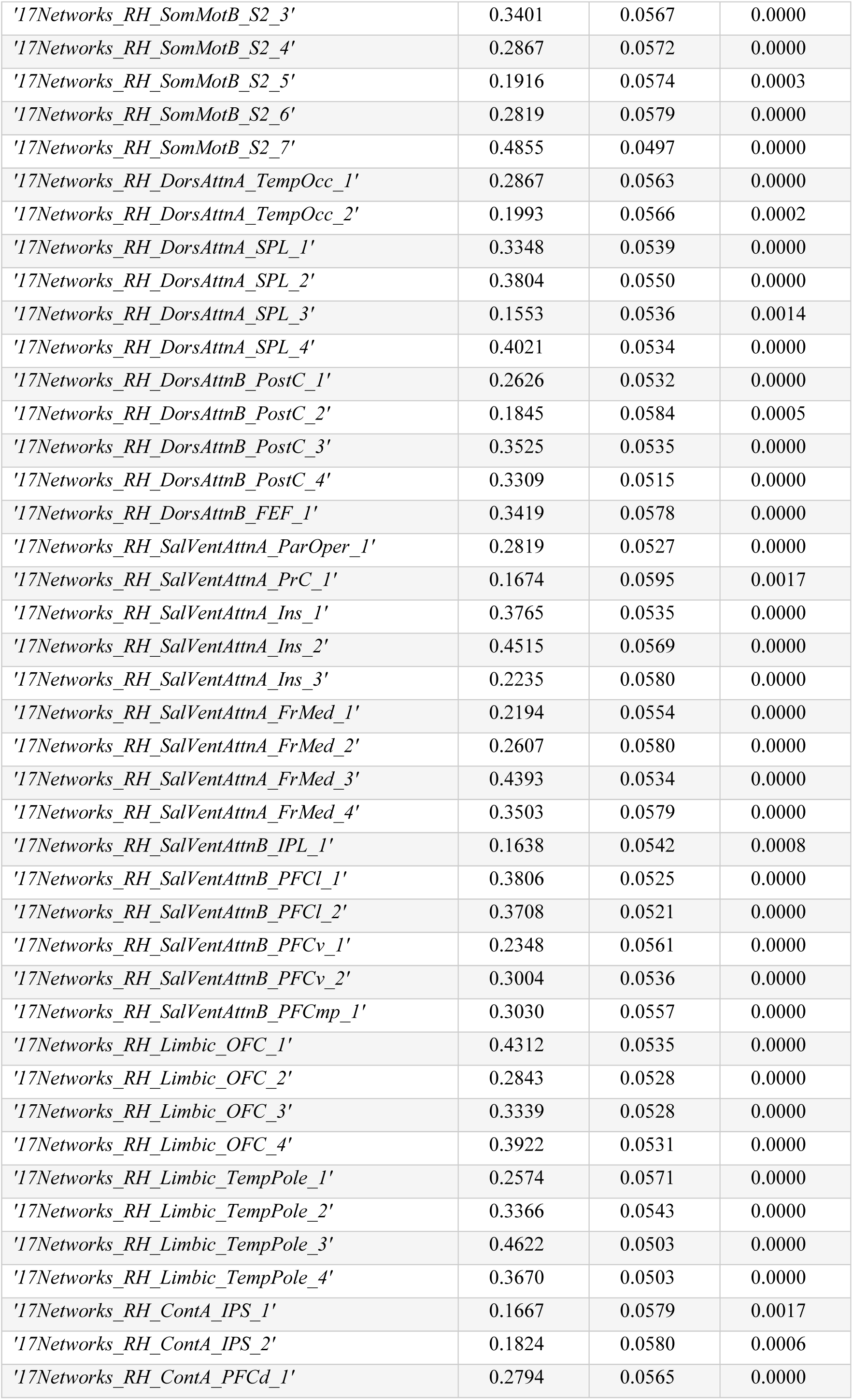
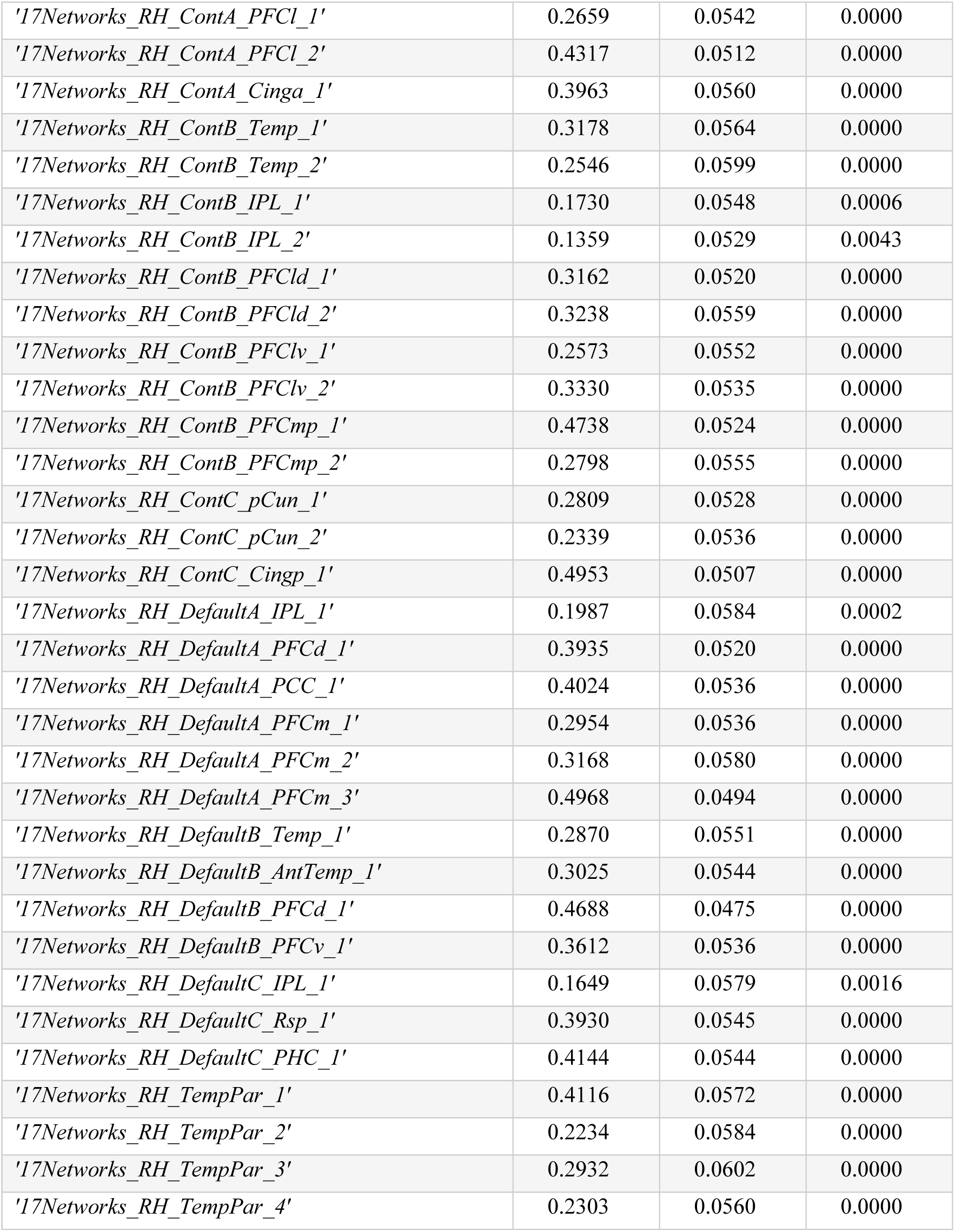
Parcel-wise heritability of cortical thickness in HCP. Heritability, standard error and p-values of cortical thickness controlling for global thickness, as well as age, sex, age*age, and age*sex using Solar 8.4.0.

**Table S8.** Post-hoc test for potential confounds (depression/BMI/intelligence) on genetic correlation between duration of sleep and cortical thickness. ** reflects FDRq<0.05, * reflects p<0.05.

| Confound | Environmental correlation | Genetic correlation |
| --- | --- | --- |
| **Depression** |  |  |
| *Left superior frontal gyrus* | 0.12 | -0.46 ** |
| *Right superior frontal gyrus* | 0.15 | -0.46 ** |
| **BMI** |  |  |
| *Left superior frontal gyrus* | 0.11 | -0.45 ** |
| *Right superior frontal gyrus* | 0.13 | -0.45 ** |
| **Intelligence** |  |  |
| *Left superior frontal gyrus* | 0.09 | -0.41 ** |
| *Right superior frontal gyrus* | 0.09 | -0.39 ** |

**Table S9.** Genetic correlations between sleep duration and cortical thickness were inconsistent across HCP and eNKI samples. Cross-sample replication of FDR-corrected ROIs from genetic correlation analysis in Fig. 2. * indicates to significant correlation (p<0.05).

| Genetic correlation HCP (complete sample) |  |  |
| --- | --- | --- |
| *Left superior frontal gyrus* | r | p |
| HCP (unrelated sample) | -0.12 | 0.014* |
| eNKI | -0.02 | 0.64 |
| HCP (total sample) | -0.07 | 0.016* |
| Genetic correlation (HCP) | -0.46 | 0.0001* |
| Environmental correlation (HCP) | 0.12 | 0.0503 |
| *Right superior frontal gyrus* | r | p |
| HCP (unrelated sample) | -0.08 | 0.11 |
| eNKI | -0.02 | 0.70 |
| HCP (total sample) | -0.06 | 0.03* |
| Genetic correlation (HCP) | -0.46 | 0.0002* |
| Environmental correlation (HCP) | 0.15 | 0.014* |

**Table S10.** Posthoc association between latent component 1 and behavior in HCP sample. The correlation between the first behavioral saliency and markers of behavior in HCP. Only correlations at FDRq<0.05 are shown.

|  |  |
| --- | --- |
| CogTotalComp_Unadj | **r=0.98** |
| CogFluidComp_Unadj | r=0.82 |
| CogCrystalComp_Unadj | r=0.76 |
| ReadEng_Unadj | r=0.72 |
| PicVocab_Unadj | r=0.68 |
| ListSort_Unadj | r=0.55 |
| CardSort_Unadj | r=0.54 |
| ProcSpeed_Unadj | r=0.54 |
| PicSeq_Unadj | r=0.50 |
| Flanker_Unadj | r=0.48 |
| SSAGA_Educ | r=0.44 |
| IWRD_TOT | r=0.29 |
| SSAGA_Income | r=0.26 |
| NEOFAC_O | r=0.23 |
| MMSE_Score | r=0.23 |
| DDisc_AUC_40K | r=0.23 |
| DDisc_AUC_200 | r=0.22 |
| LifeSatisf_Unadj | r=0.19 |
| NEOFAC_A | r=0.12 |
| AngHostil_Unadj | r=-0.11 |
| NEOFAC_N | r=-0.11 |
| PSQI_Comp5 | r=-0.12 |
| DSM_Anxi_Pct | r=-0.12 |
| DSM_Somp_Pct | r=-0.12 |
| DSM_Antis_Raw | r=-0.13 |
| SSAGA_TB_Still_Smoking | r=-0.13 |
| PSQI_Comp2 | r=-0.14 |
| PercStress_Unadj | r=-0.14 |
| PSQI_Score | r=-0.16 |
| PSQI_Comp3 | r=-0.18 |
| Weight | r=-0.25 |
| SSAGA_BMICat | r=-0.29 |
| SSAGA_TB_DSM_Difficulty_Quitting | r=-0.30 |
| BMI | r=-0.31 |

**Table S11.** Post-hoc association between latent component 2 and behavior in HCP sample. The correlation between the second behavioral saliency and markers of behavior in HCP. Only correlations at FDRq<0.05 are shown.

|  |  |
| --- | --- |
| CogTotalComp_Unadj | r=0.70 |
| CogFluidComp_Unadj | r=0.61 |
| CogCrystalComp_Unadj | r=0.52 |
| DSM_Depr_Pct | r=0.49 |
| ReadEng_Unadj | r=0.48 |
| PicVocab_Unadj | r=0.47 |
| ProcSpeed_Unadj | r=0.42 |
| Flanker_Unadj | r=0.40 |
| ListSort_Unadj | r=0.39 |
| CardSort_Unadj | r=0.38 |
| PicSeq_Unadj | r=0.34 |
| PSQI_Comp7 | r=0.31 |
| PSQI_Score | r=0.31 |
| DSM_Avoid_Pct | r=0.29 |
| DSM_Adh_Pct | r=0.28 |
| Sadness_Unadj | r=0.28 |
| AngAffect_Unadj | r=0.28 |
| FearSomat_Unadj | r=0.27 |
| Weight | r=0.26 |
| DSM_Inat_Raw | r=0.25 |
| DSM_Anxi_Pct | r=0.25 |
| BMI | r=0.25 |
| SSAGA_Depressive_Sx | r=0.24 |
| Loneliness_Unadj | r=0.24 |
| NEOFAC_N | r=0.24 |
| FearAffect_Unadj | r=0.23 |
| DSM_Hype_Raw | r=0.23 |
| SSAGA_Educ | r=0.22 |
| SSAGA_BMICat | r=0.21 |
| DSM_Antis_Raw | r=0.21 |
| NEOFAC_O | r=0.20 |
| AngHostil_Unadj | r=0.19 |
| PSQI_Comp4 | r=0.19 |
| DSM_Somp_Pct | r=0.19 |
| SSAGA_Depressive_Ep | r=0.17 |
| PercStress_Unadj | r=0.17 |
| PSQI_Comp5 | r=0.17 |
| PSQI_Comp2 | r=0.15 |
| IWRD_TOT | r=0.15 |
| MMSE_Score | r=0.14 |
| PSQI_Comp3 | r=0.11 |
| PercReject_Unadj | r=0.11 |
| PercHostil_Unadj | r=0.11 |
| SSAGA_ChildhoodConduct | r=0.11 |
| SSAGA_MOBorn | r=-0.11 |
| LifeSatisf_Unadj | r=-0.12 |
| EmotSupp_Unadj | r=-0.14 |
| Friendship_Unadj | r=-0.16 |
| NEOFAC_E | r=-0.17 |
| MeanPurp_Unadj | r=-0.18 |
| PosAffect_Unadj | r=-0.21 |
| SSAGA_TB_DSM_Difficulty_Quitting | r=-0.21 |
| NEOFAC_C | r=-0.28 |

## Supplementary Results

### Functional decoding of neurogenetic markers of sleep

To enable inference about functional brain processes that are sustained by the regions showing significant phenotypic and genetic correlation with sleep duration, we performed quantitative functional decoding using the BrainMap database^33^. Here, we observed the left superior frontal cortex, phenotypically implicated in sleep duration in the HCP unrelated sample, to be activated by tasks involving (socio-) cognitive, action, and emotion processes (p<0.05). In the eNKI sample we observed phenotypic correlation in bilateral inferior temporal regions as well as right posterior occipital cortex, which were all associated with action observation, perception, as well as spatial cognition (p<0.05). In addition, inferior temporal regions were associated with spatial cognition as well as negative affect and interoception (p<0.05). Last, the left medial frontal cortex and right superior frontal cortex, which had genetic correlation with sleep duration, were implicated in processes involving (socio)-cognition and emotion (Figure S4).

### Post-hoc association between latent behavioral saliencies and behavioral markers in HCP

Last, to further qualify the latent behavioral components, we correlated the latent behavioral components derived from the eNKI data to HCP-based markers of behavior and health (Table S10 and S11). Again, we observed strong positive relationships between component 1 and intelligence, sleep amount (Component 3, inversely coded), and negative associations with BMI, and global sleep score. Thus, component one seems to reflect a positive-negative axis of behavior, relating healthy sleep hygiene to positive behaviors whereas low sleep quality related negatively to this factor. On the other hand, component 2 reflected intelligence, depression, global sleep quality, and weight, but also negative emotions (mental) health issues, and loaded negative to friendship, extraversion and life satisfaction. Thus, component places sleep on an axis of behavior that corresponds to ruminative/worry behavior versus life satisfaction.

